# Pathogenic *MYBPC3* missense variants alter protein-protein interactions within the sarcomere

**DOI:** 10.64898/2026.05.27.727676

**Authors:** Andrea D. Thompson, Chelsea Pankiewicz, Lucy S. Plenge, Ulla Lilienthal, Sruthi Kotaru, Mathav Vignesh, Trisha Phan, Christopher McAllister, Jaime Yob, Michael P. Morley, Jodie Ingles, Sophie Hespe, Adam S. Helms, David Ginsburg, SHaRe-HCM Investigators, Sharlene M. Day

## Abstract

**Background:** Hypertrophic cardiomyopathy (HCM) is a genetic heart disease that leads to left ventricular hypertrophy, heart failure, and arrhythmias. Pathogenic missense variants in the gene myosin binding protein C (*MYBPC3)* cluster within its internal subdomains C3 and C6. The protein (MyBP-C), carrying these missense variants, localizes normally to the myofilaments, leaving uncertainty regarding the mechanism(s) by which they cause HCM.

**Methods:** We probed the molecular pathogenesis of these variants by analyzing (1) their prevalence in an international registry of patients with HCM, (2) total MyBP-C levels and the allelic fraction of mutant MyBP-C in human left ventricular heart tissue, and (3) performing flag-immunoprecipitation and proximity labeling mass spectrometry of wild-type MyBP-C and four pathogenic missense variants (Arg495Gln, Arg502Trp- C3, Trp792Arg, Arg810His- C6) to determine the change in MyBP-C interacting and proximity proteins induced by these variants.

**Results:** We found that among patients with HCM who had any *MYBPC3* pathogenic variant, 17.9% had a missense variant within the C3 or C6 subdomain. Unlike truncating variants, these C3 or C6 missense variants did not reduce MyBP-C content relative to myosin. The mutant allelic fraction of MyBP-C varied from 10.0-67.0% across samples. Flag-immunoprecipitation mass spectrometry identified 252 MyBP-C interacting proteins. Pathogenic missense variants disrupted 23 MyBP-C protein interactions, including lysosomal Ragulator-Rag complex proteins (RRAGA, RRAGC, LAMTOR4). Proximity labeling mass spectrometry was more sensitive, identifying 3,240 MyBP-C proximity proteins. Pathogenic missense variant(s) altered the proximity to MyBP-C of 789 proteins (69.4% increased and 30.5% decreased). Proteins that were increased in proximity to the missense MyBP-C were enriched for thin-filament proteins.

**Conclusion:** Pathogenic *MYBPC3* missense variants within the C3 and C6 subdomains are present in a substantial subset of patients with HCM. Our findings imply unique mechanisms of these variants distinct from haploinsufficiency, potentially driven by enhanced proximity to the thin filament within myofilaments.

## Introduction

Pathogenic variants in myosin binding protein C3 (*MYBPC3*) are the most common genetic cause of hypertrophic cardiomyopathy (HCM).^1^ Of patients with pathogenic variants in *MYBPC3*, ∼85% have truncating variants in *MYBPC3,* while the other 15% have non-truncating variants, which are mostly missense single amino acid substitutions.^2^ It is well established that truncating pathogenic variants in *MYBPC3* are evenly distributed throughout the gene and result in a premature stop codon^2^ that leads to haploinsufficiency and no detectable truncated mutant protein.^3–6^ In contrast, *MYBPC3* missense variants cluster tightly in 3 domains: C3, C6, and C10. We have shown previously that variants in the C10 domain result in mislocalization and markedly shortened protein half-lives.^2^ Conversely, variants in C3 and C6 localize normally in the A-band of myofilaments and demonstrate half-lives similar to WT MyBP-C.^2^ Therefore, the mechanisms by which they cause disease must be distinct from those of truncating variants and non-truncating variants in C10.

Structural characterization of WT MyBP-C and selected missense variants has provided some clues to their potential mechanisms. Pathogenic missense variants in the C3 subdomain cluster within a surface-exposed region, which is predicted to alter the electrostatic properties of the surface.^2,7^ However, the MyBP-C protein interactions at this molecular surface remain undefined. The C3-C6 subdomains form a flexible bridge between thin and thick filaments.^8,9^ Cryo-electron microscopy has demonstrated that the C5 subdomain binds the interacting heads motif (IHM) on myosin in the presence of the allosteric myosin inhibitor mavacamten.^8^ Biochemical studies using microscale electrophoresis of isolated fragments have also shown that the central domain of MyBP-C interacts with myosin^10,11^ and that genetic variants may disrupt this interaction. Further cryo-electron microscopy has modeled interactions of the N-terminal domains (CO-C2) of MyBP-C with the thin filament.^12,13^ Still, it remains unclear if alteration of MyBP-C binding to myosin and/or the thin filament represents a common mechanism for pathogenic variants within the central C3 and C6 subdomains.

Prior work from our laboratory used flag- immunoprecipitation mass spectrometry (IP/MS) to define WT MyBP-C-protein interactions in cardiomyocytes.^4^ In the current study, we have leveraged this technique to determine whether missense variants in C3 and C6 result in altered MyBP-C-protein interactions compared to WT MyBP-C. While this commonly employed pull-down methodology allows for the identification of protein binding partners, it is limited in its ability to distinguish differential binding to highly abundant proteins that cannot be eradicated from the control IP conditions. Because we were interested in capturing differential interactions between the flexible central domain of MyBP-C with sarcomere binding partners *in situ* that are transient and dynamic, we employed a complementary proximity labeling mass spectrometry (PL/MS) technique. We used the TurboID system, an engineered biotin ligase that labels proteins in proximity to MyBP-C (∼10-35 nm) with biotin.^14^ TurboID has faster labeling kinetics (∼10 minutes) and a higher degree of labeling than the original BioID, making it ideal for probing dynamic interactions with much higher temporal resolution. In this study, we use both IP/MS and PL/MS in cardiomyocytes to determine how pathogenic missense variants in C3 (Arg495Gln, Arg502Trp) and C6 (Trp792Arg and Arg810His) alter protein binding and the neighborhood of proteins proximal to MyBP-C, respectively.

## Methods

### SHARE analysis – *MYBPC3* variants

The generation of the Sarcomeric Human Cardiomyopathy (SHaRe) registry has been previously described.^1^ Data for this study were exported on 06/26/2025 (2025Q2). Institutional review board and ethics approval were obtained in accordance with policies applicable to each SHaRe site. Inclusion criteria included patients with a clinical diagnosis of HCM (phenotype-negative family members not included).

Similar to prior analysis,^2^ with 2025Q2 data among the patients in SHaRe who had undergone genetic testing, we identified patients with a *MYBPC3* missense variant, defined as a non-synonymous, non-truncating, single amino acid substitution (Supplemental Table 1). Variants were classified as pathogenic or likely pathogenic (P/LP), variants of unknown significance (VUS), benign or likely benign (B/LB) in accordance with American College of Medical Genetics and Genomics Association for Molecular Pathology (ACMG/AMP) standards,^15,16^ with the oversight of the SHaRe variant adjudication committee. Missense *MYBPC3* variants were reviewed for potential splice variants by cross-referencing variants with *in silico* and mini-gene assay analysis of *MYBPC3* variants predicted to impact splicing.^17–19^ Missense variants present at donor position -1 or experimentally shown to cause splicing were considered truncating

### Myectomy tissue analysis

Myocardial tissue collected at the University of Michigan or the University of Pennsylvania was obtained as previously described^5,14^ from subjects with HCM at the time of surgical myectomy to treat left ventricular outflow tract obstruction (n=21) or heart transplantation (n =2).

Collection of human myocardial tissue was performed under protocols and ethical regulations approved by Institutional Review Boards at both institutions. Patients gave informed consent. Before tissue retrieval, all hearts were perfused with ice-cold cardioplegia. All tissues were flash frozen in liquid nitrogen either immediately after myectomy (n = 21) or collected as transmural myocardial samples dissected from the mid LV free wall below the papillary muscle from failing HCM hearts at the time of transplantation (n =2).

### Alpha-LISA – measurement of MyBP-C and MYH protein levels

Alpha-LISA was performed as previously described^20^ to evaluate MyBP-C and myosin (MYH) protein levels in myectomy samples. Briefly, myectomy samples were lysed using a homogenizer in RIPA lysis and extraction buffer with protease inhibitor and phosphatase inhibitor cocktails at a concentration of 20 mg/ml. Total tissue lysate was analyzed for protein concentration using a DC protein assay, and samples were diluted to 25 μg/ml in Alpha-LISA lysis buffer (Revvity, AL003C). Five μl of this tissue lysate (25 μg/ml) from each patient sample was tested in triplicate in both the MyBP-C and MYH assays. Samples from patients with HCM who were genetically tested and were genotype negative (Sarc -) were used as negative controls, and results were normalized to these samples (100% MyBP-C, 100% MYH, MyBP-C/MYH ratio 100%). Samples from patients with HCM with a pathogenic truncating *MYBPC3* variant were utilized as a positive control, as these samples have previously been demonstrated to have a reduced MyBP-C/MYH ratio by mass spectrometry. ^5^ The ratio of normalized MyBP-C/normalized MYH*100% is reported for each sample in triplicate. Additional experimental details of this protocol can be found within the supplemental materials.

### AQUA Mass spectrometry

AQUA mass spectrometry was performed by MS Bioworks (Ann Arbor, MI). A detailed protocol is provided in the supplemental materials. Briefly, left ventricular tissue was processed as described in the supplemental material, and MyBP-C within gel bands was treated with elastase (Arg502Trp samples, Promega) at 37 °C for 18 hours and Trypsin (Trp792Arg, Arg810His, Gly531Arg samples, Promega) at 37 °C for 4 hours. The supernatant was analyzed directly without further processing. The resultant peptide solution was spiked with 200 fmol of 3 AQUA peptides per experiment (Supplemental Table 2). Peptides were analyzed in analytical duplicate by nano LC/RM using a Waters NanoAcquity HPLC system interfaced to a ThermoFisher Fusion Lumos mass spectrometer as described within the supplemental materials. The peptide m/z values used are detailed in Supplemental Table 2. Data Processing: Data were processed using Skyline v21.2. The spiked gel digests were analyzed in analytical duplicate. Peak areas were calculated using Skyline. Peak areas for duplicate injections, light/heavy ratios, average, stdev., %CV and converted fmol amounts are contained within the accompanying Supplemental Table 3.

### Cardiomyocyte Preparation and Culture

#### Isolation of Neonatal Rat cardiomyocytes (NRVMs)

NRVMs were isolated as previously described.^4^ Additional details can be found in the supplemental materials.

#### Induced pluripotent stem cell (iPSC) derived cardiomyocytes (IPSC-CM)

iPSC-CMs used in this study are from a gene-edited iPSC homozygous *MYBPC3* knockout line (*MYBPC3* [–/–]).^21^ Stem cell maintenance, cardiomyocyte production, and cell culture were performed as previously described.^20,21^ Additional details can be found in the supplemental materials.

#### Expression of *MYBPC3* via recombinant adenovirus

Turbo-Flag-*MYBPC3* and Flag- *MYBPC3* mutants were generated by site-directed mutagenesis as previously described^2^ using the QuikChange II XL Kit (Agilent) from WT human *MYBPC3* cDNA. Adenovirus was generated with the ViraPower Adenoviral Gateway Expression Kit (Invitrogen) using the pAd/CMV/V5-DEST Gateway vector and amplified in HEK293A cells.

#### Flag-MyBP-C and Turbo-Flag-MyBP-C immunofluorescence (IF)

Immunofluorescence was performed as previously described.^2,20^ Additional details of the protocol can be found in the supplemental materials. Briefly, *MYBPC3* [-/-] iPSC-CMs were plated at 150,000 cells per well in a 24-well plate (Costar #3526) on PDMS coverslips (Specialty manufacturing, Saginaw, Michigan). 4 days after replating (day 20 of differentiation), flag-*MYBPC3* or turboID-flag-*MYBPC3* pathogenic missense variants were expressed via adenoviral transduction at MOI 10, media was changed 48 hours later. At 96 hours, cells were washed with PBS and fixed for 15 minutes at room temperature in 2% paraformaldehyde (Sigma-Aldrich). Cells underwent immunofluorescence staining as described in the supplemental materials. Cells were either stained with primary and secondary antibodies (as described in the supplemental materials) to visualize flag-MyBP-C localization or with labeled streptavidin (as described in the supplemental materials) to visualize the localization of biotinylated proteins. Each sample was tested in biological duplicates, and six independent images per coverslip were acquired using a Nikon Eclipse Ti-E inverted fluorescence microscope on a 40X Plan Fluor DIC M/N2 (Nikon) objective.

#### Flag-MyBP-C immunoprecipitation (IP)

Flag-MyBP-C IP was performed as previously described.^4^ Additional details of the protocol can be found in the supplemental materials. Briefly, Flag-MyBP-C IP was performed on whole cell lysate from cells that were non-transduced (negative control), transduced with WT flag-*MYBPC3* (positive control), or C3 and C6 pathogenic missense variants flag-*MYBPC3* Arg495Gln, Arg502Trp, Trp792Arg, Arg810His, tested in biologic duplicates. 20 μl anti-FLAG M2–conjugated Sepharose beads (Sigma-Aldrich) were incubated overnight at 4 °C with 500 μg/100 μl total cellular lysate, and Flag-IP was performed as described in the supplemental materials. Flag-MyBP-C interacting proteins were eluted by competitive binding in 100μl with 100μg/ml 3X FLAG peptide (Sigma-Aldrich) for 2 hours at 4 °C. Ninety μl of the elution sample was submitted to mass spectrometry and 10 μl was utilized for flag western blot and silver stain analysis as described in more detail within the supplemental materials. LC-MS/S of flag-IP samples was performed by MS Bioworks (Ann Arbor, MI). Thirty microliters of the flag-IP (elutant samples) were separated by 10% Bis-Tris Novex mini-gel (Invitrogen) using the MES buffer system. The gel was stained with Coomassie, and each lane was excised into ten equally sized segments. Gel pieces were processed as described in the supplemental materials.

#### Protein identification of flag-IP samples using LC-MS/MS

The gel digests were analyzed by nano LC/MS/MS with a Waters M-class HPLC system interfaced to a ThermoFisher Fusion Lumos using a data-dependent acquisition (DDA) method as described in the supplemental materials. Data were filtered 1% protein and peptide level false discovery rate (FDR) and required at least two unique peptides per protein. The full list of proteins was evaluated, and known contaminants were excluded, resulting in unnormalized spectral counts. These were then converted to normalized spectral abundance factors (NSAF). Based on the equation.

NSAF = (SpC/MW)/Σ(SpC/MW)_N_
Where SpC = Spectral Counts
MW = Protein MW in kDa
N = Total Number of Proteins

#### Defining *MYBPC3* interacting proteins by LC-MS/MS

Criteria for identifying *MYBPC3* interacting proteins were the following: (a) proteins were detected in both experimental replicates for a given sample, (b) proteins showed a calculated fold-change score in normalized spectral abundance factor (NSAF) of 2 or higher over the non-transduced samples.

#### Identifying changes in MyBP-C interacting proteins in the presence of a pathogenic missense variant(s)

Criteria for proteins differentially associated with MyBP-C in the presence of pathogenic missense variants is as follows (a) proteins showed a 50% increase or decrease in NSAF compared to WT MyBP-C samples, and exhibited (b) FDR adjusted p-value (q-value) < 0.05, p-values were calculated using a two-tailed 2 -sample t-test and were corrected for multiple comparisons by calculating q-value using a two-stage step up method of Benjamini, Krieger, and Yekutieli. We limited our analysis to proteins identified as *MYBPC3-interacting* proteins. NSAF were compared for WT samples and individual pathogenic variants. We also compared all pathogenic variants as a single group (WT vs pathogenic variants-Arg495Gln, Arg502Trp, Trp792Arg, Arg810His) and pathogenic variants within a given subdomain (WT vs C3 pathogenic variants-Arg495Gln and Arg502Trp samples, WT vs C6 pathogenic variants- Trp792Arg and Arg810His variants).

#### Biotin-ligase based proximity labeling

TurboID is an engineered biotin ligase that uses ATP to convert biotin into biotin–AMP, a reactive intermediate that covalently labels proximal proteins. Optimized by directed evolution, TurboID has substantially higher activity than previously described biotin ligase–related proximity labeling methods, such as BioID, enabling higher temporal resolution and broader applications in vivo.^14^ A detailed protocol for Biotin-ligase-based proximity labeling for WT turboID-flag-MyBP-C and four pathogenic variants (Arg495Gln, Arg502Trp, Trp792Arg, Arg810His) can be found in the supplemental materials. Briefly, iPSC-CMs gene-edited to be a homozygous *MYBPC3* knockout line (*MYBPC3* [–/–])^21^ were plated at a cellular density of 2 x 10^6^ cells per well and cultured until day 20 of differentiation. Then, cells were transduced using adenoviruses at MOI 10 for 48 hours, carrying one of the following constructs: flag-*MYBPC3* – negative control, turboID-flag-*MYBPC3* WT – positive control, or a turboID-flag *MYBPC3* pathogenic mutant: Arg495Gln, Arg502Trp Trp792Arg, or Arg810His. Each sample was tested in 3 biological replicates. Four days post-infection (day 24 of differentiation), biotin labeling was performed in RPMI + P/S with 50 μM biotin (Sigma B4501) and 500 μM ATP (Sigma) prepared freshly. After 1 hour of biotin labeling, cells were washed 3 times with PBS (Gibco) and lysed by adding 500 μl/well of lysis buffer as described in the supplemental materials. A streptavidin-Biotin pull-down was performed using 100 μg of total cell lysate per sample, incubated overnight at 4°C with 750 μl of Dynabeads MyOne streptavidin C1 beads (Invitrogen, prewashed three times with lysis buffer within Lobind 1.5 ml microcentrifuge tubes). Each sample’s beads were washed as described within the supplemental materials, and then 650 μl of beads were submitted for mass spectrometry to identify biotinylated proximity proteins bound to streptavidin beads. LC-MS/MS was performed by MS Bioworks (Ann Arbor, MI). On-bead trypsin digestion was performed manually with the protocol described in detail within the supplemental materials. Peptides were taken to dryness using a lyophilizer. Peptides were reconstituted in 0.1% TFA for mass spectrometry analysis. 100 μl was removed for western blot and silver stain analysis (elutant) as described in the supplemental materials.

#### Protein identification of MyBP-C PL samples using LC-MS/MS data-dependent acquisition (DDA)

Ten percent of each sample was analyzed in analytical triplicate by LC/MS with a ThermoFisher Vanquish Neo UPLC system interfaced to a ThermoFisher Orbitrap Astral. The mass spectrometer was operated in data-dependent acquisition mode. Additional details of this protocol can be found in the supplemental materials. The full list of proteins was reported as unnormalized spectral counts (SpC). These were then converted to normalized spectral abundance factors (NSAF). Based on the equation.

NSAF = (SpC/MW)/Σ(SpC/MW)_N_
Where SpC = Spectral Counts
MW = Protein MW in kDa
N = Total Number of Proteins

#### Defining MyBP-C proximity proteins by LC-MS/MS

Criteria for identifying *MYBPC3* proximity proteins were the following: (a) proteins were detected in all experimental and technical replicates, and (b) proteins showed a calculated fold-change score in normalized spectral abundance factor (NSAF) of 2 or higher over the negative control samples or were absent in the negative controls. To create volcano plots, data were processed using log2-transformed intensity values.

#### Changes in MyBP-C proximity, using LC-MS/MS data-independent acquisition (DIA)

We utilized data-independent acquisition (DIA) mass spectrometry to identify changes in MyBP-C proximity proteins between WT MyBP-C and individual MyBP-C carrying pathogenic. This is described in detail within the supplemental materials. Briefly, ten percent of each sample was analyzed in analytical triplicate by LC/MS with a ThermoFisher Vanquish Neo UPLC system interfaced to a ThermoFisher Orbitrap Astral. The mass spectrometer was operated in data-independent mode as described in detail within the supplemental materials. DIA data were analyzed suing DIA-NN (v.1.9.1) which severed several functions: conversion of RAW files to QUANT, alignment based on retention times, searching data using *in-silico* spectral library created from the FASTA file and iteratively using the spectral library created from RAW data, filtering of database search results at 1% peptide and protein false discovery rate (FDR), calculation of peak areas for detected peptides, data normalization. The report.pg_matrix.tsv output was uploaded to Perseus 1.5.5.3 for further analysis. Proteins were reported as normalized protein intensity. Data was processed log2 transforming intensity values, with missing values replaced with 0.

#### Identifying proteins with altered MyBP-C proximity in the presence of a pathogenic missense variant(s)

Differences in normalized intensity values were compared for proteins present in all WT and pathogenic variant samples and were identified by performing a student’s 2-tailed 2-sample t-test. P-values were corrected for multiple comparisons by calculating q-value with Benjamini-Hochberg FDR correction (FDR 5%). Proteins with altered proximity were defined as proteins with a fold change of log2 transformed values > 0.5 or < -0.5 (increase or decrease in *MYBPC3* binding) and FDR-adjusted p-value (q-value) < 0.05.

### Gene Ontology Enrichment Analysis

Gene ontology analysis was performed using Metascape [http://metascape.org] ^22^ and TOPPGENE ^23^ and is described in more detail in the supplemental materials.

## Results

### Prevalence of pathogenic *MYBPC3* missense variants in patients with HCM in the SHaRe registry

At the time of data extraction from SHaRe,^1^ 8,496 patients with HCM underwent genetic testing, of whom 2,656 patients had a variant of any type in *MYBPC3.* Of these, 1,707 patients had pathogenic truncating variants, 148 had in-frame duplications and deletions, and 811 patients had non-synonymous, non-truncating missense variants. Of the non-synonymous, non-truncating missense variants, 291 were unique variants (Supplemental Table 1). Most of these variants were classified as variants of uncertain significance or benign. Only 24 of the 291 missense variants were adjudicated as pathogenic/likely pathogenic (P/LP), identified in 305 patients. P/LP missense variants are present in 17.9% of all patients with a *MYBPC3* P/LP variant (305/1,707) and 3.6% of all patients with HCM who underwent genetic testing (305/8,496) (Supplemental Table 1). In 83.6% of patients who had a P/LP *MYBPC3* missense variant (255/303), the variant was located in the C3 or C6 subdomain (Figure 1A, Supplemental Table 1).

**Figure 1:**
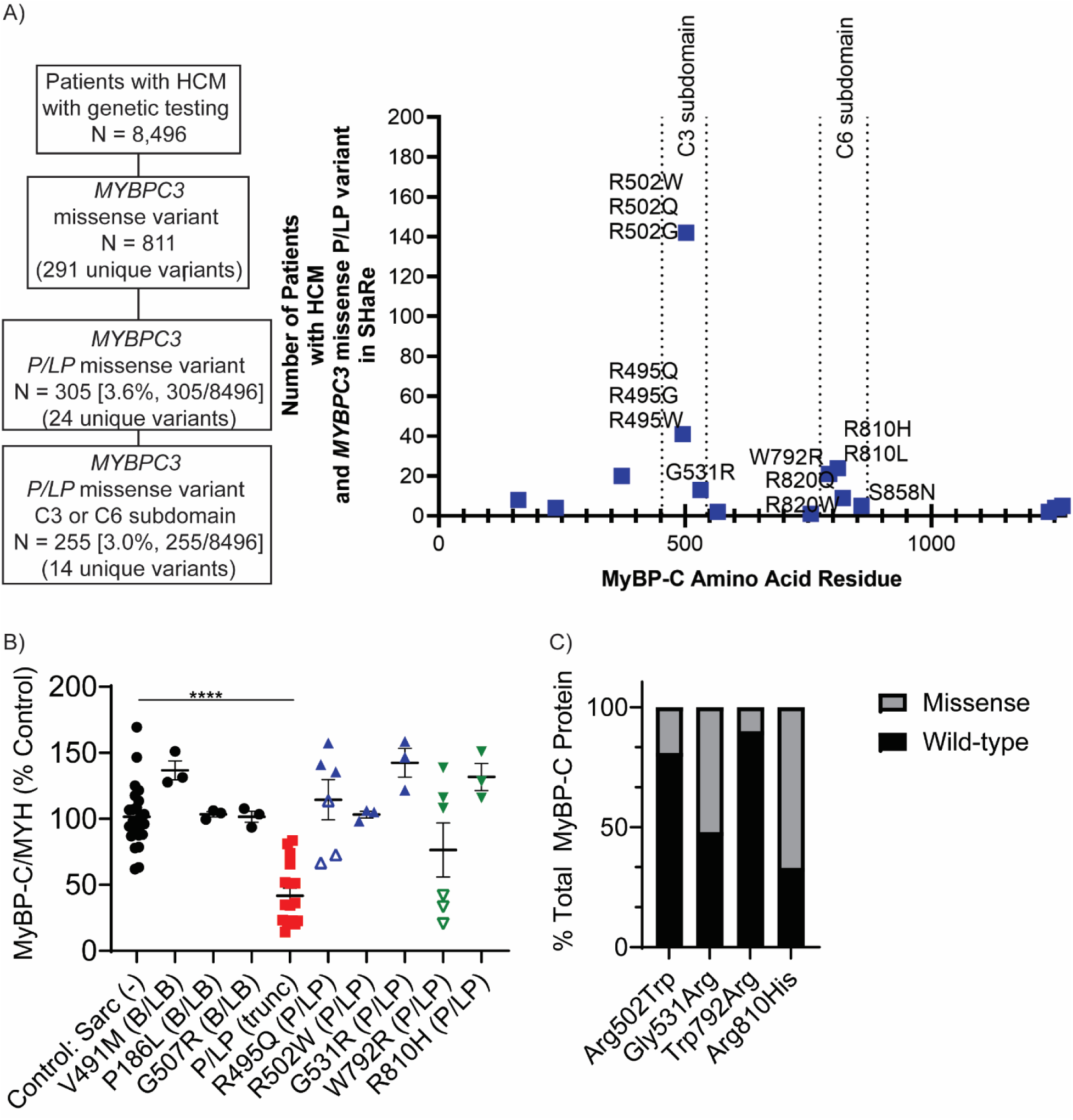
Prevalence and consequences of P/LP *MYBPC3* missense variants in C3 and C6. **A)** Prevalence of pathogenic/likely pathogenic (P/LP) missense variants in SHaRe registry is 3.6% of all patients with HCM and genetic testing. P/LP *MYBPC3* missense variants cluster in the C3 and C6 subdomains. 86.3% of HCM patients with a P/LP *MYBPC3* missense variant have a variant within these two subdomains. The single-letter amino acid code is utilized in this figure. **B)** The ratio of MyBP-C/MYH was measured in triplicate by alpha-LISA for each patient sample. Left ventricular tissue from patients with HCM and *MYBPC3* P/LP truncating variants (red squares) has a reduced level of MyBP-C/MYH ratio compared to genotype negative patients with HCM or those with benign variants [black circles] (p-value <0.0001, ****). P/LP missense variants are shown in blue upright triangles for the C3 subdomain and green downward triangles for the C6 subdomain exhibit no significant difference in MyBP-C/MYH ratio compared to controls. R495Q and W792R had two patient samples available for testing, and samples from individual patient samples are indicated by open and closed triangles. Single letter amino acid code is utilized in this figure. **C)** Considerable variability is observed in the % of protein expression with the missense variant, ranging from 10% of total protein Trp792Arg in one patient sample and 67% of total protein for the missense variant Arg810His.

### Most pathogenic *MYBPC3* missense variants do not cause haploinsufficiency and result in variable mutant protein expression

Consistent with our prior work using mass spectrometry ^5,6^, *MYBPC3* truncating variants result in a reduced amount of total MyBP-C protein relative to myosin (MyBP-C/MYH-mean 43.25% +/- standard deviation 25.14%) compared to genotype-negative patients (Sarc-) (MyBP-C/MYH- 100% +/- 23.86%, p-value < 0.0001) or patients with B/LB variants (Figure 1B). Conversely, P/LP *MYBPC3* missense variants in the C3 subdomain (Arg495Gln, Arg502Trp, Gly531Arg) or C6 subdomain (Trp792Arg, Arg810His) do not alter MyBP-C protein content (Figure 1B), except for a single patient with Trp792Arg who demonstrated a decreased MyBP-C/MYH ratio.

We performed AQUA-MS on LV tissue from four patients, each with a single P/LP *MYBPC3* missense variant, as described previously for Arg495Gln.^17^ The percentage of mutant MyBP-C varied considerably from 10% to 74.4% in patient samples (Figure 1C). The fraction of mutant protein shows a linear positive correlation with cellular half-lives previously measured^2^ (Supplemental Figure 1, R^2^ = 0.90), consistent with protein stability being the primary determinant of the fraction of mutant protein in the heart.

### Flag immunoprecipitation identifies 252 MyBP-C interacting proteins. We first defined MyBP-C

interacting proteins by performing flag-IP DDA mass spectrometry for N-terminal flag-tagged human MyBP-C (WT) and P/LP *MYBPC3* missense variants in C3 (Arg495Gln, Arg502Trp) or C6 (Trp792Arg, Arg810His) expressed in NRVCMs. Using non-transduced NRVCM lysate as a negative control, we identified 252 MyBP-C interacting proteins (Supplemental Table 4). As expected, the bait (human MyBP-C) was the most abundant protein. Rat MyBP-C was identified as an interacting protein of human MyBP-C. There was significant overlap among interacting proteins between WT and P/LP variant samples (Supplemental Figure 2A). Gene ontology analysis of interacting proteins shows that MyBP-C interacting proteins are enriched in myofibril proteins in all samples (Supplemental Figure 2B).

### Flag-immunoprecipitation identifies MyBP-C protein interactions that are lost in the presence of pathogenic *MYBPC3* missense variants

Protein input in cellular lysate samples for flag-IP was equivalent between samples (Figure 2A). There were no significant differences in MyBP-C interacting proteins for individual P/LP C3 variants compared to WT MyBP-C (Figure 2B, Supplemental Table 5). In contrast, several interacting proteins exhibited decreased binding to C6 mutant MyBP-C (Supplemental Table 5, Figure 2B) compared to WT MyBP-C. Of the 30 proteins identified to have decreased binding for either Trp792Arg or Arg810His, 19 disrupted binding in both samples (Figure 2D). Proteins with decreased binding to C6 mutant protein were enriched in proteins outside of the sarcomere (Figure 2C) such as mitochondria protein containing complex (lsca2, Nsf1, Tomm40), mitochondria-associated ER membrane contact site (Tomm40, Tmx1, Rab32), and GTPase activity (Rab8a, Rab21, Atl3, Rac1, Rab32). Only 2 interacting proteins were increased for the Trp792Arg mutant protein compared to WT: Cox7b (mitochondrial inner membrane) and Hmgb1 (nuclease).

**Figure 2.**
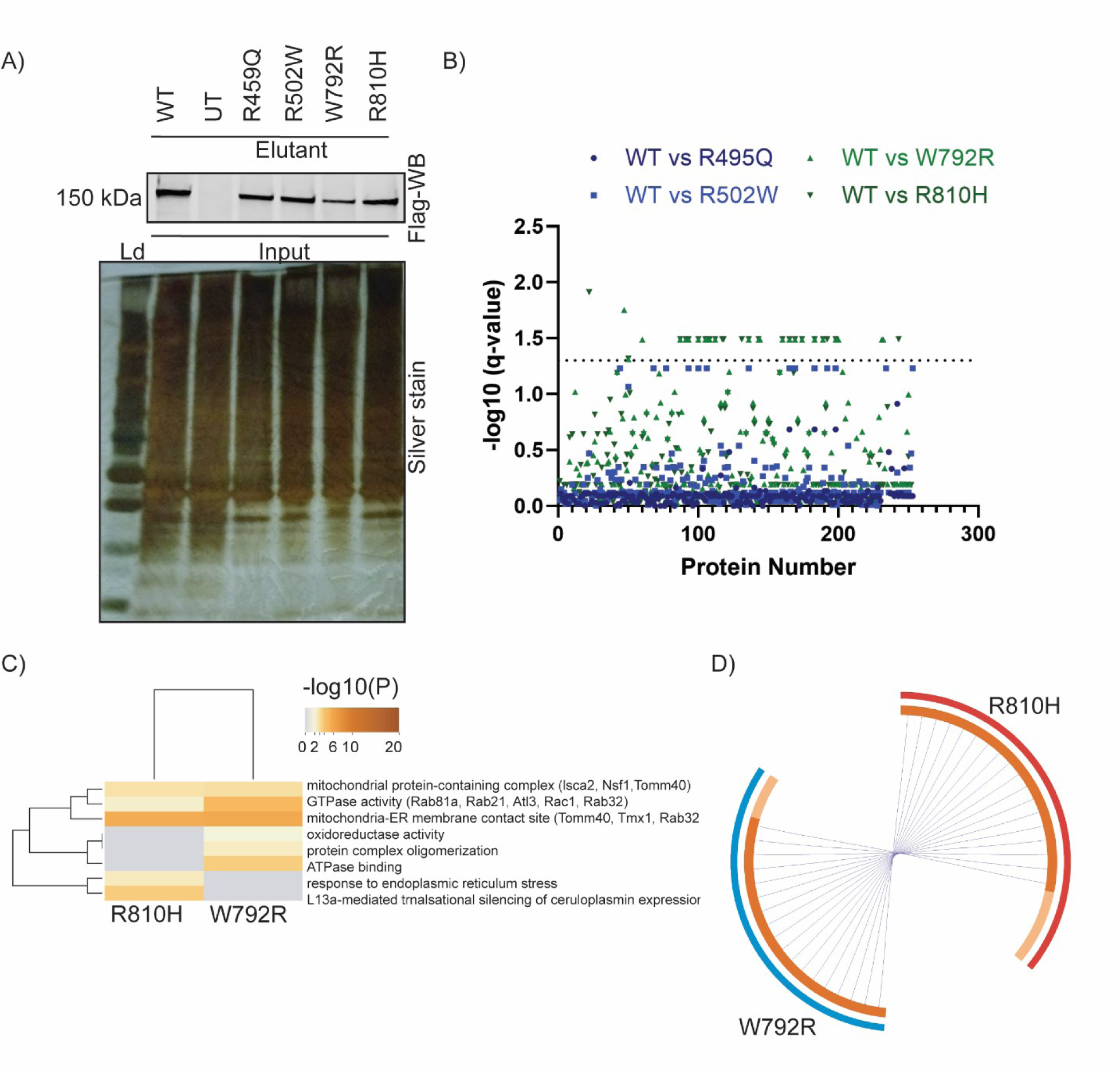
Comparison of Flag-MyBP-C interacting proteins between wild-type and mutant missense MyBP-C. A) Flag-MyBP-C immunoprecipitation (IP) gel analysis. After cellular lysis (input), flag immunoprecipitation was performed, and elutant samples were submitted for mass spectrometry with two biological replicates performed per condition. Gel analysis was performed on one biological replicate per sample for quality control. There is equal protein input in cellular lysate samples that underwent flag-IP (silver stain). Flag western blot analysis of the elutant identified the presence of bait protein (flag-MyBP-C). Conditions included Flag-MyBP-C wild-type (WT), Arg495Gln (R495Q), Arg502Trp (R502W), Trp79Arg (W792R), Arg810His (R810H). Flag-MyBP-C is absent in the non-transduced negative control (UT). **B) Differentially associated MyBP-C interacting proteins.** A Manhattan plot (protein number of x-axis, -log10 q-value on y-axis) visualizes proteins identified to have increased or decreased binding to MyBP-C. Individual pathogenic variants are as follows: Arg495Gln (R495Q, dark blue circle), Arg502Trp (R502W, blue squares), Trp792Arg (W792R, upward green triangle), Arg810His (R810H, downward green triangle). **C) Gene ontology enrichment analysis of differentially associated MyBP-C interacting proteins.** Using metascape, we identified all statistically enriched terms and selected the term with the best p-value within each cluster as its representative term and displayed them in a dendrogram. The dendogram cells are colored by their p-values; grey cells indicate the lack of enrichment for that term in the corresponding gene list. Proteins with decreased binding to MyBP-C Trp792Arg (W792R) and Arg810His (R810H) compared to WT are enriched in proteins outside the sarcomere. Specifically, GO:0098798, GO:00003924, GO:0044233, GO:0016491, GO:0051259, GO:0051117, GO:0034976, R-RNO-156827). **D) Comparison of differentially associated protein partners for MyBP-C Trp792Arg and Arg810His**. The Circos plot shows how genes from the input gene lists of proteins with decreased binding compared to WT overlap. On the outside, each arc represents the identity of each gene list (Arg810His (R810H)-red, Trp792Arg (W792R)-blue). On the inside, each arc represents a gene list, where each gene member of that list is assigned a spot on the arc. Dark orange color represents the genes that are shared by both lists and light orange color represents genes that are unique to that gene list. Purple lines link the genes that are shared by both gene lists. 19/30 proteins (63%) that exhibited decreased binding to either MyBP-C Trp792Arg and/or Arg810His compared to WT were the same in both samples.

To increase statistical power, we pooled data from all 4 mutant proteins and compared them to WT MyBP-C. This analysis identified 23 proteins that exhibited decreased interactions for mutant vs. WT MyBP-C (Figure 3A, Supplemental Table 6). No proteins exhibited increased binding to pathogenic missense proteins. Gene ontology analysis of these 23 proteins revealed enrichment for proteins within the GTPase activator complex, specifically LAMTOR4, RRAGA, and RRAGC(Figure 3B). The protein-protein interaction network MCODE algorithm shows that LAMTOR4-RRAGC-RRAGA are in the same protein complex (Figure 3C). This Ragulator-Rag mTORC1 activation complex is located on the surface of lysosomes.^24^

**Figure 3.**
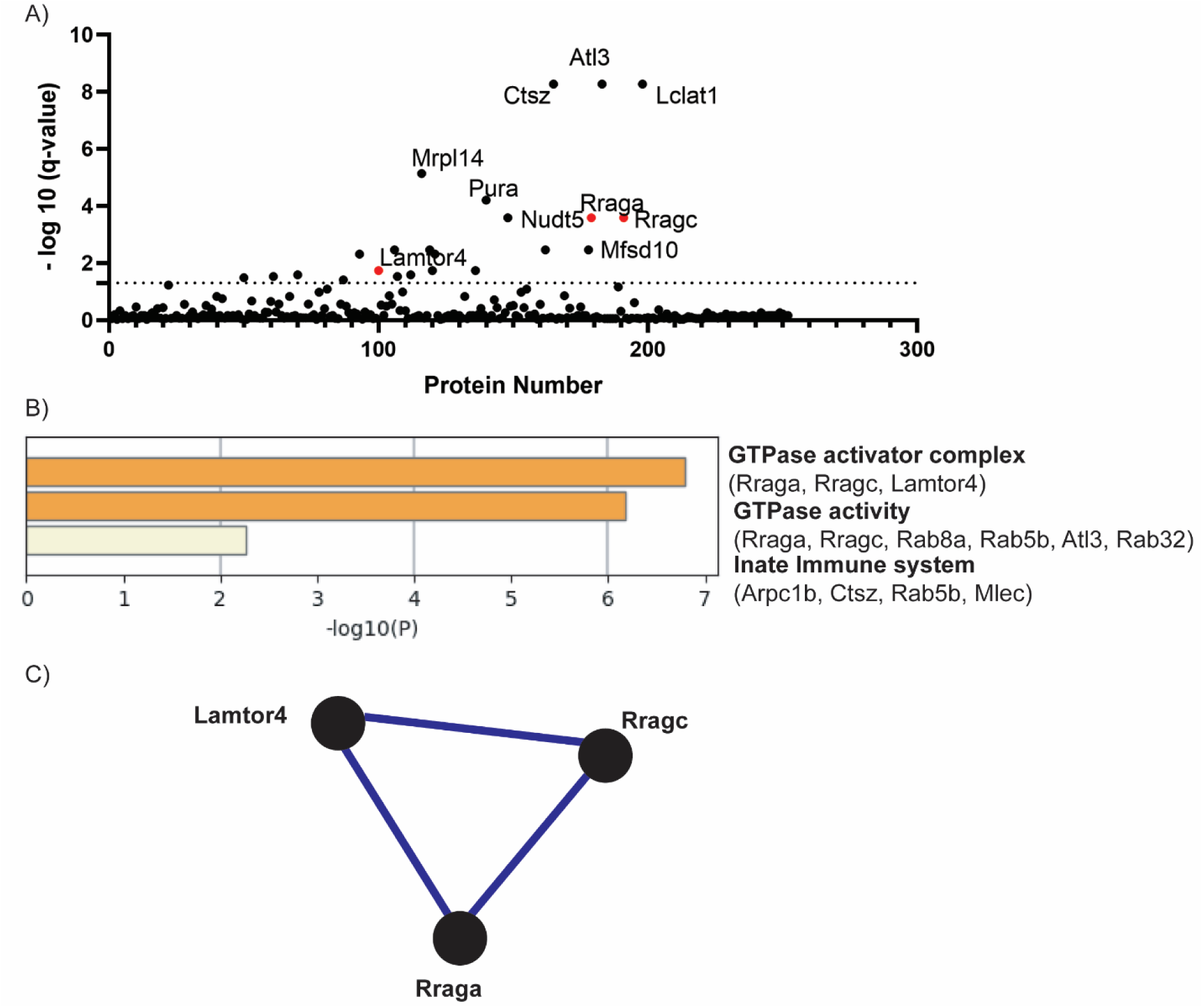
Differential MyBP-C-protein interactions pooling all mutant protein samples compared to WT MyBP-C. A) Differentially associated MyBP-C interacting proteins. A Manhattan plot (protein number of x-axis, -log10 q-value on y-axis) visualizes proteins identified to have increased or decreased binding to MyBP-C carrying a P/LP missense variant. In this analysis, Arg495Gln, Arg502Trp, Trp792Arg, and Arg810His samples were compared to WT. All proteins that were differentially associated with MyBP-C demonstrated decreased binding in the P/LP missense variant sample (Supplemental Table 6) RRAGA, RRAGC, and LAMTOR4 as red circles. **B) Gene ontology enrichment of proteins with decreased interactions for mutant proteins.** Using metascape, we identified all statistically enriched terms and selected the term with the best p-value within each cluster as its representative term and displayed them in a bar graph (-log 10 p-value). The most enriched pathway was GTPase activator complex (GO: 19027773) with RRAGA, RRAGC, and LAMTOR4 having reduced binding to MyBP-C carrying P/LP missense variants. Also enriched are GTPase activity (GO:0003924) and innate immune system (R-RNO-168249). **C) Decreased binding of mutant MyBP-C to the Ragulator-Rag complex proteins (RRAGA, RRAGC, LAMTOR4) by M-CODE algorithm.** Gene enrichment analysis applied to each MCODE network to extract “biological meanings”. The top three p-values were identified as the following gene ontology terms. GO:1990877 (FNIP-folliculin RagC/D GAPI), GO:1902773 (GTPase activator complex), and GO:0150005 (enzyme activator complex). RRAGA-RRAGC-LAMTOR4 are known as binding partners, illustrated by MCODE network.

### Proximity labeling identifies the neighborhood of MyBP-C proximity proteins

While we identified loss of protein binding partners for mutant compared to WT MyBP-C, we were unable to resolve differential association with highly abundant proteins (e.g. sarcomere) that cannot be eradicated from control IP conditions. We also suspected that we were not capturing transient and dynamic interactions between the flexible central domain of MyBP-C with sarcomere binding partners. To overcome these limitations, we employed a complementary proximity labeling mass spectrometry (PL/MS) technique to define whether the C3 and C6 mutants altered the protein neighborhood of MyBP-C.

We performed proximity labeling using TURBO-ID by transducing WT or mutant proteins via adenoviral vectors into iPSC-CMs gene-edited to be a homozygous *MYBPC3* knockout line (*MYBPC3* [–/–]).^21^ Proximity proteins were identified by DDA mass spectrometry (Supplemental Table 7) and DIA mass spectrometry (Supplemental Table 8). Like flag-MyBP-C, Turbo-Flag-MyBP-C proteins localized normally to myofibrils (Supplemental Figure 3). Biotin labeling of proteins was also localized within myofibrils (Supplemental Figure 4). One biologic replicate of Arg810His had a low level of protein detection in both DIA and DDA mass spectrometry, comparable to the negative control (flag-MyBP-C), suggesting failure of on-bead trypsinization. This sample was excluded from further analysis.

We identified MyBP-C proximity proteins using DDA mass spectrometry (Figure 4A) and compared results to flag immunoprecipitation DDA mass spectrometry. Proximity labeling identified a much larger number of proteins proximal to MyBP-C (n=3,240) compared to interacting proteins by IP/MS (n=252). The number of proximal proteins was 1,823 for WT, 1,078 for Arg495Gln, 2,826 for Arg502Trp, 2,897 for Trp792Arg, and 2,965 for Arg810His (Supplemental Table 9, Supplemental Figure 5). There was significant overlap between the groups, with a total of 964 proteins identified in all samples. Among the most abundant 500 proximity proteins, gene ontology showed enrichment for myofibril proteins across all WT and mutant samples (Supplemental Figure 5-7). Given this increased sensitivity, we were interested in further exploring the cellular compartment and molecular function of WT MyBP-C proximity proteins. Using a second gene ontology software TOPPGENE,^23^ we found that the most significant enrichment in molecular function among WT MyBP-C proximity proteins was cytoskeletal protein binding (Supplemental Figure 6) and that the most significant enrichment in cellular component was in myofibrils and cytoplasm (Supplemental Figure 7). This is consistent with the known molecular function and cellular localization of wild-type MyBP-C. Of the 252 proteins identified as MyBP-C interacting proteins by IP/MS, only 99 (39.3%) were also identified as proximity proteins by PL/MS (Supplemental Table 10). RRAGA and RRAGC were among those proteins identified as both MyBP-C-interacting proteins and MyBP-C-proximity proteins (Supplemental Table 10).

**Figure 4:**
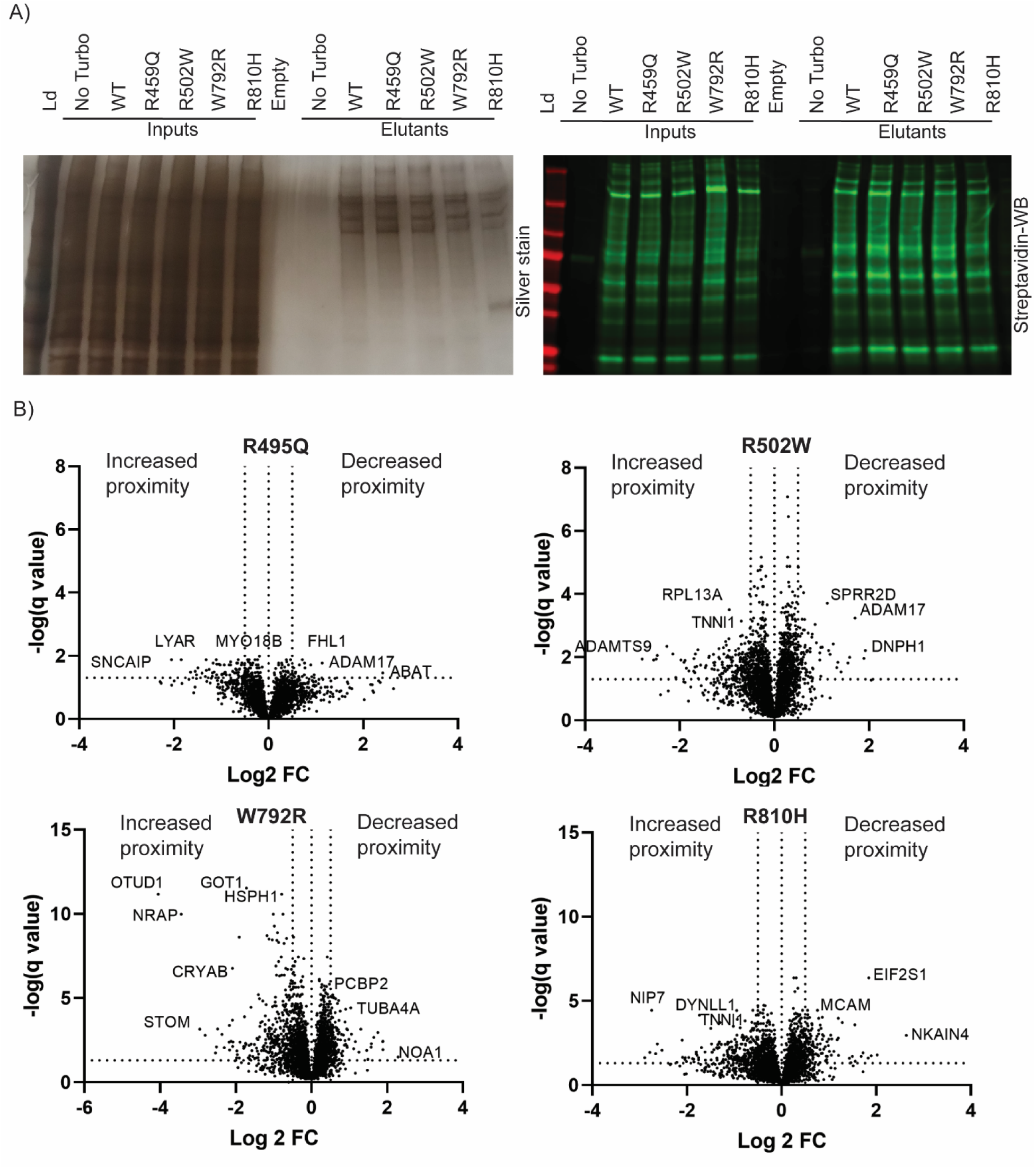
MyBP-C proximity proteins with increased or decreased proximity to MyBP-C relative to WT. **A) Proximity Labeling Gel Analysis.** After proximity labeling, the cellular lysate (input) was subjected to streptavidin pull-down, and elutant samples were submitted to mass spectrometry for DDA and DIA mass spectrometry. Three biological replicates were analyzed in technical triplicate per sample by mass spectrometry. One biological replicate was analyzed by gel analysis per sample for quality control. Total protein was visualized by silver stain, and biotinylated proteins are visualized by streptavidin western blot. Ld = ladder, No turbo = flag-MyBP-C without TurboID biotin ligase as a negative control. Conditions included TurboID-Flag-MyBP-C wild-type (WT), Arg495Gln (R495Q), Arg502Trp (R502W), Trp79Arg (W792R), Arg810His (R810H). **B) Increased or decreased proximity relative to WT MyBP-C.** Volcano plots [x-axis: log_2_ fold change (FC) and y-axis: -log_10_ (q-value)] comparing DIA mass spectrometry results visualize proteins with increased and decreased proximity to individual pathogenic mutant MyBP-C compared to wild-type.

### Proximity labeling mass spectrometry identifies MyBP-C proximity proteins that increase and decrease proximity to MyBP-C in the presence of a pathogenic *MYBPC3* variant

We used proximity labeling DIA mass spectrometry to identify changes in proximity proteins induced by individual missense proteins compared to WT MyBP-C (Figure 4). Altogether, 789 proteins were differentially abundant compared to WT in at least one of the mutant MyBP-C proteins, with 30.5% being decreased in abundance and 69.4% increased in abundance in mutant vs WT MyBP-C. Like IP/MS, Arg495Gln differed the least from WT MyBP-C (Figure 4,5), with only 20 proteins exhibiting decreased proximity and 55 proteins exhibiting increased proximity compared to WT MyBP-C. The other 3 mutant proteins had far more differentially abundant proximal proteins (Figure 4, Supplemental Table 11): Arg502Trp (81 decreased, 206 increased), Trp792Arg (124 decreased, 383 increased), and Arg810His (140 decreased, 265 increased). There was significant overlap between proteins with decreased (Figure 5A) or increased proximity (Figure 5B) across these three variants. Proximal proteins RAGA and RRAGC were neither decreased nor increased in abundance in WT vs. mutant proteins (Supplemental Table 11).

**Figure 5:**
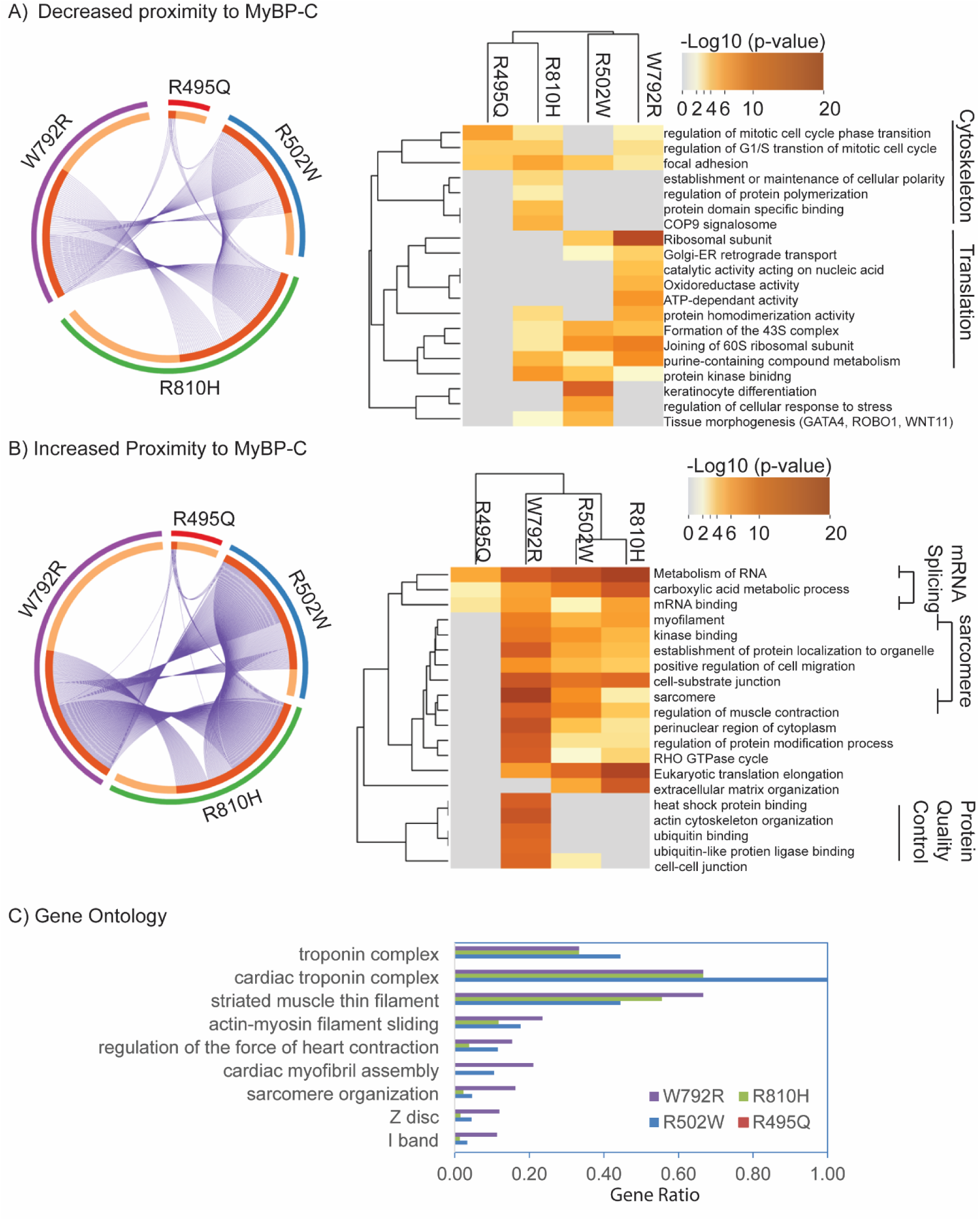
Proximity proteins that exhibit altered proximity to the missense MyBP-C compared to WT MyBP-C. Using metascape, we performed gene ontology enrichment analysis and a circos plot and dendogram are provided, similar to those described in figure 2 for **(A) Proteins with decreased proximity to a missense MyBP-C compared to WT, and (B) Proteins with increased proximity to a missense MyBP-C compared to WT.** The Circos plot shows how genes from the input gene lists of proteins with decreased MyBP-C proximity compared to WT (Panel A) or increased MyBP-C proximity compared to WT (Panel B) overlap. On the outside, each arc represents the identity of each gene list (Arg495Gln -R495Q, red, Arg502Trp – R502W, blue, Trp792Arg – W792R, purple, Arg810His- R810H, green). On the inside, each arc represents a gene list, where each gene member of that list is assigned a spot on the arc. Dark orange color represents the genes that are shared by multiple lists, and light orange color represents genes that are unique to that gene list. Purple lines link the same gene that are shared by multiple gene lists. Again Arg495Gln (R495Q) has the fewest proteins decreased (Panel A) or increased (Panel B) in proximity compared to WT MyBP-C. For the other 3 mutant proteins, there is significant overlap in protein identity (Circos plot). Using Metascape, we identified all statistically enriched terms and selected the term with the best p-value within each cluster as its representative term and displayed them in a dendrogram. The dendogram cells are colored by their p-values; grey cells indicate the lack of enrichment for that term in the corresponding gene list. Proteins with decreased proximity of mutant compared to WT MyBP-C (Panel A) are enriched in proteins involved in translation (GO:0044391 ribosomal subunit and R-HAS-72695 Formation of 43S complex) and cytoskeletal structure (GO:0005925 focal adhesion and GO:1901990 regulation of mitotic cell cycle phase transition). Proteins with increased proximity to MyBP-C compared to WT MyBP-C (Panel B) are enriched in mRNA binding (R-HAS-8953854 Metabolism of RNA, GO:0003729 mRNA binding) and myofibrils (GO:0036379 myofilament, GO: 0030017 sarcomere). Some pathways are uniquely enriched in specific variants, for example, MyBP-C Trp792Arg is enriched in pathways involved in protein quality control (GO:0031072 heat shock protein binding, GO:0043130 ubiquitin binding). **C) Subcategorization of proteins within the myofilament compartment with increased proximity for mutants compared to WT MyBP-C.** We subcategorized the enriched terms within the clusters represented by myofilament, sarcomere, and regulation of muscle contraction and graphed the gene enrichment ratio (# gene ID for identified in proteins with increased proximity/ # gene ID within the term) for each of these terms. For Arg502Trp (R502W), Trp792Arg (W792R), and Arg810His (R810H) MyBP-C, the most enriched proteins within these categories are the cardiac troponin complex (GO:1990584).

By gene ontology analysis, missense mutant proteins exhibit decreased proximity for proteins involved in focal adhesion, the cytoskeleton, the mitotic cell cycle, and protein translation (Figure 5A). Two proteins (ADAM17 and EIF3J) had decreased proximity in all mutant MyBP-C proteins, and 32 proteins had decreased proximity in the presence of 3 MyBP-C pathogenic variants (Supplemental Table 12). Only 2 proteins overlapped between PL/MS and IP/MS as being decreased in the mutant vs. the WT protein. EIF2S1 had decreased interaction with MyBP-C Trp792Arg and Arg810His (Supplemental Table 5) and decreased proximity to MyBP-C Arg502Trp, Trp792Arg, and Arg810His (Supplemental Table 11). The lysosomal hydrolase Nudt5 had decreased interaction with all 4 MyBP-C mutants compared to WT (Supplemental Table 6) and decreased proximity to MyBP-C Arg502Trp, Trp792Arg, and Arg810His (Supplemental Table 11).

In contrast to proteins with decreased proximity to MyBP-C being largely outside of the sarcomere, proteins with increased proximity to MyBP-C Arg502Trp, Trp792Arg, and Arg810His were enriched within the sarcomere (Figure 5B). This enrichment was driven specifically by increased proximity to proteins within the striated muscle thin filament (TNNC1, TNNI1, TNNI3, TNNT2, TPM1, TPM4, TMOD3, LMOD2) and cardiac troponin complex (TNNC1, TNNI3, TNNT2) (Figure 5C). One hundred and three proteins exhibited increased proximity to MyBP-C in the presence of 3 or more of the pathogenic variants tested, including TNNI1, TNNT2, and TNNC1 (Supplemental Table 12).

Increased proximity of MyBP-C to proteins involved in mRNA binding and regulation was observed across all 4 P/LP *MYBPC3* missense variants. The summary terms metabolism of RNA and mRNA binding are driven by the more specific ontology terms of the spliceosome and regulation of alternative mRNA splicing. Finally, there were unique findings for individual pathogenic variants. For example, MyBP-C Trp792Arg had increased proximity proteins involved in protein quality control, such as heat shock protein binding and ubiquitin binding.

## Discussion

This work provides new insights into the prevalence and unique mechanisms of pathogenic missense variants in *MYBPC3*. First, we leverage a growing population in the international SHaRe registry to establish the prevalence of pathogenic missense variants in patients with HCM, comprising 17.9% of all patients with a pathogenic variant in *MYBPC3* and 3.6% of all genotyped patients. The vast majority (86.3%) of pathogenic missense variants are within the subdomains C3 and C6. Next, we demonstrated that, in contrast to truncating variants, pathogenic missense variants do not result in haploinsufficiency in total MyBP-C protein within human HCM hearts. Allelic protein imbalance was a common finding, with mutant protein accounting for 10-74% of total protein across samples. The positive correlation of mutant protein fraction with protein half-lives, published previously, suggests that the inherent stability of the mutant proteins is a primary determinant of the content of mutant protein relative to wild-type protein in the sarcomere. This concept is further supported by the consistency of the mutant allele fraction for the same pathogenic variant in unrelated patients (i.e. Arg495Gln – 62% and 69%).^17^

With haploinsufficiency excluded as a mechanism for missense *MYBPC3* variants, we hypothesized that altered protein-protein interactions or altered positioning of the protein within the sarcomere underlie their pathogenic properties. To address this, we employed 2 complementary techniques to identify proteins that interact with MyBP-C (IP/MS) or are in proximity (PL/MS). Each of these techniques has strengths and limitations. While IP/MS identifies proteins that are binding partners, abundant proteins often contaminate the control IP samples in the absence of an antibody, potentially obscuring differential binding to a wild-type vs mutant protein. Another limitation of IP/MS is that the cells are lysed prior to pull-down, mixing proteins from different cellular compartments that would not normally interact in the intact cell, potentially resulting in false positive interactions. PL/MS overcomes the inherent limitations of IP/MS, but cannot distinguish whether a proximal protein is in the vicinity (i.e. within the “biotin cloud”) or is a binding partner.

In comparing results from these 2 techniques, we identified both overlapping and distinct findings. PL/MS identified ∼13-fold higher number of proximity proteins compared to interacting proteins identified with IP/MS. This is not surprising, as interacting proteins would be expected to be a subset of proteins in proximity to MyBP-C. However, only 39% of interacting proteins from IP/MS were validated with PL/MS, suggesting that perhaps many of the remaining interacting proteins may be false positives, i.e. are not in the same vicinity in the intact myocyte. In contrast, proximal proteins to WT MyBP-C identified by PL/MS were enriched within the myofilament compartment, including contractile proteins and proteins involved in RNA regulation, translation, proteostasis and metabolism that have been identified previously with proximity labeling of *ACTN2* and *TTN*.^25,26^ This result suggests this technique is better suited to define the proximity of sarcomere proteins and conformational changes within the sarcomere.

Of those proteins that did validate as both interacting and proximal proteins, RRAGA and RRAGC were of particular interest as they have not been shown previously to bind to MyBP-C. They are members of the Ragulator-Rag mTORC1 activation complex located on the surface of lysosomes. In the presence of amino acids, Ragulator (LAMTOR1-5) scaffolds and binds an active heterodimer Rag GTPase complex (GTP bound RagA/B and GDP bound RagC/D) which recruits mTORC1 to the surface of the lysosome for activation.^24^ mTORC1 activation is known to drive cardiac hypertrophy and can be attenuated by rapamycin, an inhibitor of mTORC1.^27,28^ mTORC1 can also be inactivated by protein binding to the Ragulator-Rag complex.^29^ By IP/MS, the interaction between WT MyBP-C and the Ragulator-Rag complex proteins was absent for all missense mutant proteins, suggesting disrupted binding. However, the MyBP-C proximity was not different between WT and missense mutant MyBP-C proteins by PL/MS, suggesting disrupted binding but preserved localization in the presence of missense variants. Further investigation is needed to validate this novel MyBP-C interacting partner and explore if disruption of this interaction by pathogenic missense variants may lead to MTORC1 activation and left ventricular hypertrophy.

Overall, there was minimal overlap in the differential abundance of interacting proteins by IP/MS with proximity proteins by PL/MS between the missense mutant and wild-type MyBP-C. By IP/MS, only 30 proteins were decreased or absent in at least one of the two C6 mutants, and no interacting proteins were different between the C3 mutants compared to WT MyBP-C. When the mutant samples were pooled to increase statistical power, only 23 interacting proteins were differentially abundant in the mutants, and all of them exhibited decreased or absent binding. This contrasts with 789 proximity proteins identified as differentially abundant between mutant and WT MyBP-C, with most (69.4%) displaying increased proximity. We therefore conclude that proximity proteomics displayed much higher sensitivity to IP/MS in detecting potential gain-of-function shifts in the position of missense MyBP-C within the sarcomere and changes in MyBP-C protein environment.

The most striking finding from PL/MS was a marked increase in abundance of thin filament proteins for 3 of the 4 missense proteins (Arg502Trp, Trp792Arg, and Arg810His) compared to WT MyBP-C. MyBP-C is a flexible, elongated protein ∼40-44 nm in length.^30^ It is oriented in a transverse orientation with the N-terminus (where TURBO-ID was fused), transiently interacting with and activating the thin filament in a calcium and phosphorylation-dependent manner. Proximity labeling identified proteins within 10-35 nm from the N-terminus of MyBP-C as it toggles back and forth between the thick and thin filaments throughout the cardiac cycle.^5,31^ The increased abundance of thin filament proteins in missense mutant MyBP-C suggests that these pathogenic variants in the C3 and C6 domains are causing a conformational change in the protein that allosterically favors interactions of the N-terminus (and/or C3-C6) with the thin filament. This would be expected to cause activation of the thin filament, increased calcium sensitivity, and higher diastolic tension. The precise molecular interaction between the MyBP-C and the thin filament has not been fully elucidated.^13^ Further structural characterization is needed to understand how these mutant proteins are potentially altering this interaction. Further, additional work is needed to study the effect of these missense variants on cross-bridge kinetics and the biochemical and structural state of myosin.

RNA-binding proteins involved in splicing, RNA stability, and regulation were the only proteins increased in proximity in all 4 mutant proteins compared to WT MyBP-C. The identification of proteins involved in RNA regulation and protein translation at the sarcomere has uncovered an emerging role of mRNA trafficking and local translation at the Z-disc of the sarcomere.^32,33^ Whether increased proximity of missense mutant proteins to RNA machinery is indicative of altered local RNA regulation or protein translation, and whether specific transcripts are affected, will require further studies to elucidate.

The pathogenic variant Trp792Arg displayed some unique proximity protein alterations compared to the other mutants. Specifically, there was a reduced abundance of proteins involved in translation and an increased abundance of proteins involved in proteostasis, including HSP70 chaperones (HSPA8, HSPA1A), co-chaperones (DNAJB2, DNAJC6, DNAJC7, DNAJC16, BAG4), the co-chaperone E3 ubiquitin ligases (STUB1), which triage proteins for degradation. This mutant protein also has the lowest mutant allele fraction (10%) and was associated with haploinsufficiency in 1 of 2 patients studied. Although the Trp792Arg mutant protein incorporates normally into the myofilament, these findings together suggest that it has reduced intrinsic stability compared to WT MyBP-C.

### Limitations

Methodological limitations of IP/MS and PL/MS are discussed above. We studied 4 pathogenic variants in C3 and C6 in which there were both common and distinct findings in proximity and interacting proteins. Although these 4 variants are the most prevalent, the results may not be representative of all variants in these and other neighboring domains. These results are hypothesis-generating for deciphering the mechanisms by which missense variants in MyBP-C cause HCM. Additional biophysical, structural, and physiological studies are needed to understand their pathophysiology fully. Prior experiments on protein fragments demonstrated high affinity between the central domains of MyBP-C and myosin S1, with C2-C4 and C5-C7 competing for binding to the same site. Further, myosin binding was increased by C6 R820Q.^10^ In our experiments, we did not detect changes in MyBP-C myosin interactions in *MYBPC3* missense proteins, despite increased interactions with the thin filament. This may be due to the tethering of the C-terminus of MyBP-C, which is required for myofilament localization. Other structural methods may be more amenable to evaluating the effect of these variants on the myosin-MyBP-C interaction.

### Conclusion

The findings from this study highlight the prevalence and clustering of *MYBPC3* missense variants in C3 and C6 and support a predominant gain-of-function mechanism characterized by increased protein-protein interactions within the sarcomere, specifically with the thin filament and proteins involved in RNA regulation. Future studies will be needed to elucidate the effects of these strengthened interactions on cross-bridge kinetics, local transcription/translation, and sarcomere proteostasis. Distinguishing unique mechanisms of pathogenic variants in *MYBPC3* may also have therapeutic implications as we progress into the era of customized and gene-targeted therapies.

## Supporting information

SupplementalTable4_FlagIP_IP

SupplementalTable6_DA_groupIP

SupplementalTable9_PP_DDA

SupplementalTable10_comparison_IP_PP_DDA

SupplementalTable11_DP_DIA

SupplementalTable12_comparison_DA_DP

## Acknowledgements

We are grateful to Dr. Samantha Harris for providing custom MyBP-C antibodies used in the alpha-LISA. We are also grateful to Dr. Richard Jones and the team at MS Bioworks for performing the mass spectrometry as described above.

## Notes

SHaRe-HCM Investigators include Carolyn Ho and Nel Lakdawala (Cardiovascular Division, Brigham and Women’s Hospital, Boston, MA, USA), Sharlene M. Day and Anjali Owens (Penn Center for Inherited Cardiovascular, Disease, Hospital of the University of Pennsylvania and the Perelman School of Medicine at the University of Pennsylvania, Philadelphia, PA, USA), Adam Helms and Sara Saberi (Department of Internal Medicine, Division of Cardiovascular Medicine, University of Michigan, Ann Arbor, MI, USA), Mark Russell (Department of Pediatrics, Division of Cardiology, University of Michigan, Annor, MI, USA), Rachel Lampert and John C. Stendahl (Department of Medicine, Section of Cardiovascular Medicine, Yale School of Medicine, New Haven, CT, USA), Vicki Parikh and Euan Ashley (Center for Inherited Cardiovascular Disease, Division of Cardiovascular Medicine, Stanford University School of Medicine, Stanford, California, USA), Jodie Ingles, Belinda Gray (Garvan Institute of Medical Research, and UNSW Sydney, Sydney NSW, Australia), Iacopo Olivotto (Cardiomyopathy Unit, Careggi University Hospital, Florence, Italy and Cardiology Unit, Meyer Children’s Hospital IRCCS, Florence, Italy), Michelle Michels, Peter Paul Zwetsloot (Department of Cardiology, Thoraxcenter, Erasmus MC Rotterdam, The Netherlands), James Ware (National Heart and Lung Institute and MRC London Institute of Medical Sciences, Imperial College London, London, UK and and Royal Brompton & Harefield Hospitals, Harefield, UK Guy’s and St. Thomas’, London UK), Lia Crotti (Department of Cardiology, San Luca Hospital, Cardiomyopathy Unit, Center for Cardiac Arrhythmias of Genetic Origin and Laboratory of Cardiovascular Genetics, Istituto Auxologico Italiano, Milan, Italy), Henning Bungaard and Anna Axelsson Raja (Department of Cardiology, Rigshospitalet University Hospital Copenhagen, Copenhagen, Denmark), Alex Pereria (Brazil), Thomas D. Ryan and Erin M. Miller (Department of Pediatrics, University of Cincinnati College of Medicine, Heart Institute, Cincinnati Children’s Hospital Medical Center, Cincinnati, OH, USA), Joseph Rossano and Kimberly Y. Lin (Division of Cardiology, Children’s Hospital of Philadelphia, University of Pennsylvania, Philadelphia, PA, USA), Dominic Abrams (Center for Cardiovascular Genetics, Boston Children’s Hospital, MA, USA), Niccolo Maurizi (Cardiomyopathy Unit, University of Florence, Florence, Italy and Service of Cardiology, University Hospital of Lausanne, Lausanne, Switzerland), Alessia Argiro (Cardiomyopathy Unit, University of Florence, Florence, Italy), Brian Claggett (Department of Biostatistics, Harvard T.H. Chan School of Public Health, Boston, MA, USA).

## References

1. Ho CY, Day SM, Ashley EA, Michels M, Pereira AC, Jacoby D, Cirino AL, Fox JC, Lakdawala NK, Ware JS, et al. Genotype and Lifetime Burden of Disease in Hypertrophic Cardiomyopathy: Insights from the Sarcomeric Human Cardiomyopathy Registry (SHaRe). Circulation. 2018;138:1387–1398. doi: 10.1161/CIRCULATIONAHA.117.033200

2. Helms AS, Thompson AD, Glazier AA, Hafeez N, Kabani S, Rodriguez J, Yob JM, Woolcock H, Mazzarotto F, Lakdawala NK, et al. Spatial and Functional Distribution of MYBPC3 Pathogenic Variants and Clinical Outcomes in Patients With Hypertrophic Cardiomyopathy. Circ Genom Precis Med. 2020;13:396–405. doi: 10.1161/CIRCGEN.120.002929

3. Vignier N, Schlossarek S, Fraysse B, Mearini G, Kramer E, Pointu H, Mougenot N, Guiard J, Reimer R, Hohenberg H, et al. Nonsense-mediated mRNA decay and ubiquitin-proteasome system regulate cardiac myosin-binding protein C mutant levels in cardiomyopathic mice. Circ Res. 2009;105:239–248. doi: 10.1161/CIRCRESAHA.109.201251

4. Glazier AA, Hafeez N, Mellacheruvu D, Basrur V, Nesvizhskii AI, Lee LM, Shao H, Tang V, Yob JM, Gestwicki JE, et al. HSC70 is a chaperone for wild-type and mutant cardiac myosin binding protein C. JCI Insight. 2018;3. doi: 10.1172/jci.insight.99319

5. O’Leary TS, Snyder J, Sadayappan S, Day SM, Previs MJ. MYBPC3 truncation mutations enhance actomyosin contractile mechanics in human hypertrophic cardiomyopathy. J Mol Cell Cardiol. 2019;127:165–173. doi: 10.1016/j.yjmcc.2018.12.003

6. Marston S, Copeland O, Jacques A, Livesey K, Tsang V, McKenna WJ, Jalilzadeh S, Carballo S, Redwood C, Watkins H. Evidence from human myectomy samples that MYBPC3 mutations cause hypertrophic cardiomyopathy through haploinsufficiency. Circ Res. 2009;105:219–222. doi:10.1161/CIRCRESAHA.109.202440

7. Zhang XL, De S, McIntosh LP, Paetzel M. Structural characterization of the C3 domain of cardiac myosin binding protein C and its hypertrophic cardiomyopathy-related R502W mutant. Biochemistry. 2014;53:5332–5342. doi: 10.1021/bi500784g

8. Dutta D, Nguyen V, Campbell KS, Padron R, Craig R. Cryo-EM structure of the human cardiac myosin filament. Nature. 2023;623:853–862. doi: 10.1038/s41586-023-06691-4

9. Tamborrini D, Wang Z, Wagner T, Tacke S, Stabrin M, Grange M, Kho AL, Rees M, Bennett P, Gautel M, et al. Structure of the native myosin filament in the relaxed cardiac sarcomere. Nature. 2023;623:863–871. doi: 10.1038/s41586-023-06690-5

10. Ponnam S, Kampourakis T. Microscale thermophoresis suggests a new model of regulation of cardiac myosin function via interaction with cardiac myosin-binding protein C. J Biol Chem. 2022;298:101485. doi: 10.1016/j.jbc.2021.101485

11. Flavigny J, Robert P, Camelin JC, Schwartz K, Carrier L, Berrebi-Bertrand I. Biomolecular interactions between human recombinant beta-MyHC and cMyBP-Cs implicated in familial hypertrophic cardiomyopathy. Cardiovasc Res. 2003;60:388–396. doi:10.1016/j.cardiores.2003.07.001

12. Risi CM, Villanueva E, Belknap B, Sadler RL, Harris SP, White HD, Galkin VE. Cryo-Electron Microscopy Reveals Cardiac Myosin Binding Protein-C M-Domain Interactions with the Thin Filament. J Mol Biol. 2022;434:167879. doi: 10.1016/j.jmb.2022.167879

13. Risi C, Belknap B, Forgacs-Lonart E, Harris SP, Schroder GF, White HD, Galkin VE. N-Terminal Domains of Cardiac Myosin Binding Protein C Cooperatively Activate the Thin Filament. Structure. 2018;26:1604–1611 e1604. doi: 10.1016/j.str.2018.08.007

14. Cho KF, Branon TC, Udeshi ND, Myers SA, Carr SA, Ting AY. Proximity labeling in mammalian cells with TurboID and split-TurboID. Nat Protoc. 2020;15:3971–3999. doi: 10.1038/s41596-020-0399-0

15. Richards S, Aziz N, Bale S, Bick D, Das S, Gastier-Foster J, Grody WW, Hegde M, Lyon E, Spector E, et al. Standards and guidelines for the interpretation of sequence variants: a joint consensus recommendation of the American College of Medical Genetics and Genomics and the Association for Molecular Pathology. Genet Med. 2015;17:405–424. doi: 10.1038/gim.2015.30

16. Kelly MA, Caleshu C, Morales A, Buchan J, Wolf Z, Harrison SM, Cook S, Dillon MW, Garcia J, Haverfield E, et al. Adaptation and validation of the ACMG/AMP variant classification framework for MYH7-associated inherited cardiomyopathies: recommendations by ClinGen’s Inherited Cardiomyopathy Expert Panel. Genet Med. 2018;20:351–359. doi: 10.1038/gim.2017.218

17. Helms AS, Davis FM, Coleman D, Bartolone SN, Glazier AA, Pagani F, Yob JM, Sadayappan S, Pedersen E, Lyons R, et al. Sarcomere mutation-specific expression patterns in human hypertrophic cardiomyopathy. Circ Cardiovasc Genet. 2014;7:434–443. doi:10.1161/CIRCGENETICS.113.000448

18. Singer ES, Ingles J, Semsarian C, Bagnall RD. Key Value of RNA Analysis of MYBPC3 Splice-Site Variants in Hypertrophic Cardiomyopathy. Circ Genom Precis Med. 2019;12:e002368. doi: 10.1161/CIRCGEN.118.002368

19. Ito K, Patel PN, Gorham JM, McDonough B, DePalma SR, Adler EE, Lam L, MacRae CA, Mohiuddin SM, Fatkin D, et al. Identification of pathogenic gene mutations in LMNA and MYBPC3 that alter RNA splicing. Proc Natl Acad Sci U S A. 2017;114:7689–7694. doi: 10.1073/pnas.1707741114

20. Thompson AD, Wagner MJ, Rodriguez J, Malhotra A, Vander Roest S, Lilienthal U, Shao H, Vignesh M, Weber K, Yob JM, et al. An Unbiased Screen Identified the Hsp70-BAG3 Complex as a Regulator of Myosin-Binding Protein C3. JACC Basic Transl Sci. 2023;8:1198–1211. doi:10.1016/j.jacbts.2023.04.009

21. Helms AS, Tang VT, O’Leary TS, Friedline S, Wauchope M, Arora A, Wasserman AH, Smith ED, Lee LM, Wen XW, et al. Effects of MYBPC3 loss-of-function mutations preceding hypertrophic cardiomyopathy. JCI Insight. 2020;5. doi: 10.1172/jci.insight.133782

22. Zhou Y, Zhou B, Pache L, Chang M, Khodabakhshi AH, Tanaseichuk O, Benner C, Chanda SK. Metascape provides a biologist-oriented resource for the analysis of systems-level datasets. Nat Commun. 2019;10:1523. doi: 10.1038/s41467-019-09234-6

23. Chen J, Bardes EE, Aronow BJ, Jegga AG. ToppGene Suite for gene list enrichment analysis and candidate gene prioritization. Nucleic Acids Res. 2009;37:W305–311. doi: 10.1093/nar/gkp427

24. Su MY, Morris KL, Kim DJ, Fu Y, Lawrence R, Stjepanovic G, Zoncu R, Hurley JH. Hybrid Structure of the RagA/C-Ragulator mTORC1 Activation Complex. Mol Cell. 2017;68:835–846 e833. doi: 10.1016/j.molcel.2017.10.016

25. Ladha FA, Thakar K, Pettinato AM, Legere N, Ghahremani S, Cohn R, Romano R, Meredith E, Chen YS, Hinson JT. Actinin BioID reveals sarcomere crosstalk with oxidative metabolism through interactions with IGF2BP2. Cell Rep. 2021;36:109512. doi: 10.1016/j.celrep.2021.109512

26. Rudolph F, Fink C, Huttemeister J, Kirchner M, Radke MH, Lopez Carballo J, Wagner E, Kohl T, Lehnart SE, Mertins P, et al. Deconstructing sarcomeric structure-function relations in titin-BioID knock-in mice. Nat Commun. 2020;11:3133. doi: 10.1038/s41467-020-16929-8

27. Shioi T, McMullen JR, Tarnavski O, Converso K, Sherwood MC, Manning WJ, Izumo S. Rapamycin attenuates load-induced cardiac hypertrophy in mice. Circulation. 2003;107:1664–1670. doi: 10.1161/01.CIR.0000057979.36322.88

28. Kaplan JL, Rivas VN, Walker AL, Grubb L, Farrell A, Fitzgerald S, Kennedy S, Jauregui CE, Crofton AE, McLaughlin C, et al. Delayed-release rapamycin halts progression of left ventricular hypertrophy in subclinical feline hypertrophic cardiomyopathy: results of the RAPACAT trial. J Am Vet Med Assoc. 2023;261:1628–1637. doi: 10.2460/javma.23.04.0187

29. Tsun ZY, Bar-Peled L, Chantranupong L, Zoncu R, Wang T, Kim C, Spooner E, Sabatini DM. The folliculin tumor suppressor is a GAP for the RagC/D GTPases that signal amino acid levels to mTORC1. Mol Cell. 2013;52:495–505. doi: 10.1016/j.molcel.2013.09.016

30. Hartzell HC, Sale WS. Structure of C protein purified from cardiac muscle. J Cell Biol. 1985;100:208–215. doi: 10.1083/jcb.100.1.208

31. Wong FL, Bunch TA, Lepak VC, Steedman AL, Colson BA. Cardiac myosin-binding protein C N-terminal interactions with myosin and actin filaments: Opposite effects of phosphorylation and M-domain mutations. J Mol Cell Cardiol. 2024;186:125–137. doi: 10.1016/j.yjmcc.2023.11.010

32. Haddad R, Sadeh O, Ziv T, Erlich I, Haimovich-Caspi L, Shemesh A, van der Velden J, Kehat I. Localized translation and sarcomere maintenance requires ribosomal protein SA in mice. J Clin Invest. 2024;134. doi: 10.1172/JCI174527

33. Scarborough EA, Uchida K, Vogel M, Erlitzki N, Iyer M, Phyo SA, Bogush A, Kehat I, Prosser BL. Microtubules orchestrate local translation to enable cardiac growth. Nat Commun. 2021;12:1547. doi: 10.1038/s41467-021-21685-4

