## Supplemental Information for "Pathogenic *MYBPC3* missense variants alter protein-protein interactions within the sarcomere"

**Table of Contents:**

1. Supplemental Methods. Pages 2-9
2. Supplemental Table 1: MYBPC3 non-synonymous missense variants identified in patients with HCM who underwent genetic testing within the ShaRe registry (2025Q2), file information Page 10
3. Supplemental Table 2: AQUA Peptides. Page 10
4. Supplemental Table 3: Peptide m/z values used in AQUA mass spectrometry. Page 11
5. Supplemental Table 4: Flag-immunoprecipitation MyBP-C interacting proteins, file information. Page 12
6. Supplemental Table 5: Differentially Associated (DA) Flag-MyBP-C interacting proteins individual variants, file information. Page 12
7. Supplemental Table 6: Differentially Associated (DA) Flag-MyBP-C, comparing grouped pathogenic variants to WT MyBP-C. file information. Page 12
8. Supplemental Table 7: Proximity labeling mass spectrometry DDA overview of protein totals. Page 13
9. Supplemental Table 8: Proximity labeling mass spectrometry DIA overview of protein totals. Page 14
10. Supplemental Table 9: TurboID proximity labeling and DDA mass spectrometry identifies MyBP-C proximity proteins. File information. Page 15
11. Supplemental Table 10: comparison of Flag-IP MyBP-C interacting proteins (IP, 252) and Proximity labeling MyBP-C proximity proteins (PP 3,240) results DDA mass spectrometry, file information. Page 15
12. Supplemental Table 11: TurboID proximity labeling and DIA mass spectrometry identifies MyBP-C proximity proteins with decreased or increased proximity to MyBP-C in the presence of P/LP *MYBPC3* missense variant, file information. Page 15
13. Supplemental Table 12: Comparison of differential MyBP-C association and differential MyBP-C proximity, file information. Page 15
14. Supplemental Figure 1: Linear correlation between previously measured cellular half-lives and % mutant protein measured by AQUA in human left ventricular tissue. Page 16
15. Supplemental Figure 2: Flag-immunoprecipitation MyBP-C interacting proteins. Page 17
16. Supplemental Figure 3: Immunofluorescence of MyBP-C protein constructs in MYBPC3 [-/-] iPSC-CMs. Page 18
17. Supplemental Figure 4: TurboID-flag-MyBP-C protein constructs result in biotinylated proteins within the sarcomere. Page 19
18. Supplemental Figure 5: MyBP-C proximity proteins identified by DDA mass spectrometry. Page 20
19. Supplemental Figure 6: WT MyBP-C Proximity proteins identified by DDA mass spectrometry molecular function gene ontology enrichment analysis. Page 21

20. Supplemental Figure 7:WT MyBP-C Proximity proteins identified by DDA mass spectrometry cellular compartment gene ontology enrichment analysis. Page 22
21. References Page 23

**Supplemental Methods:**

**AQUA Mass spectrometry:** AQUA mass spectrometry was performed on left ventricular tissue lysate by MS Bioworks (Ann Arbor, MI). Gel bands were washed with 25 mM ammonium bicarbonate, followed by acetonitrile and reduced with 10 mM dithiothreitol at 60°C followed by alkylation with 50 mM iodoacetamide at room temperature. Gel bands were then digested with elastase (Arg502Trp samples, Promega) at 37 °C for 18 hours and Trypsin (Trp792Arg, Arg810His Gly531Arg samples, Promega) at 37 °C for 4 hours; the supernatant was analyzed directly without further processing. The resultant peptide solution was spiked with 200 fmol of 3 AQUA peptides per experiment (Supplemental Table 2). Peptides were analyzed in analytical duplicate by nano LC/RM using a Waters NanoAcquity HPLC system interfaced to a ThermoFisher Fusion Lumos mass spectrometer. Fifty percent of each sample was spiked with 100 fmol AQUA mix, loaded on a trapping column, and eluted over a 75µm analytical column at 350 nL/min; both columns were packed with Luna C18 resin (Phenomenex). A 60 min gradient was employed. The mass spectrometer was operated in PRM mode without scheduling; instrument settings included 15,000 FWHM resolution, NCE 30, AGC target value 1e5, and maximum IT of 100ms. The peptide m/z values used are detailed in Supplemental Table 2. Data Processing Data were processed using Skyline v21.2. The spiked gel digests were analyzed in analytical duplicate. Peak areas were calculated using Skyline. Peak areas for duplicate injections, light/heavy ratios, average, std. dev., %CV and converted fmol amounts are contained within the accompanying Supplemental Table 3.

**Alpha-LISA – measurement of MyBP-C and MYH protein levels:** Left ventricular tissues were lysed using a homogenizer (omni tissue homogenizer TH 115V, Revvity) in RIPA lysis and extraction buffer (Thermo Fisher), protease inhibitor cocktail (complete Mini, Roche) and phosphatase inhibitor cocktail (PhosSTOP, Roche) at a concentration of 20 mg/ml. Total tissue lysate was analyzed for protein concentration using a DC protein assay, and samples were diluted to 25 µg/ml in Alpha-LISA lysis buffer (Revvity, AL003C). Five µl of this tissue lysate (25 µg/ml) was transferred per well to an alpha-LISA assay plate (384 AlphaPlate, Revvity). Total tissue lysate from each patient sample was tested in triplicate in both the MyBP-C and MYH assays. Next, 20 µl of alpha-LISA master mix was added per well. The alpha-LISA master mix for the MyBP-C assay was a 1:2,000 dilution of mouse monoclonal MyBP-C antibody (C0, Santa Cruz, sc-137180), a 1:2,000 dilution of rabbit monoclonal MyBP-C antibody (C5-C7, Samantha Harris, University of Arizona), a 1:200 dilution of anti-mouse IgG donor bead (Revvity, AS104D), and a 1:200 dilution of anti-rabbit IgG acceptor bead (Revvity AL104C) dissolved in 1X Alpha-LISA assay buffer (Revvity, AL000F). The alpha-LISA master mix for the MYH assay was a 1:2,000 dilution of mouse monoclonal αMYH antibody (abcam, ab50967), a 1:2,000 dilution of rabbit polyclonal βMYH antibody (abcam, ab228353), a 1:200 dilution of anti-mouse IgG donor bead (Revvity, AS104D), and a 1:200 dilution of anti-rabbit IgG acceptor bead (Revvity, AL104C) dissolved in 1X Alpha-LISA assay buffer (Revvity, AL000F). Plates were covered with foil seal protectant and incubated for 48 hours at room temperature. Plates were read by laser irradiation of donor beads at 680 nm, generating chemiluminescent emission from the acceptor bead at 615 nm using EnVision Multimode Plate Reader (Revvity). Samples from patients with HCM who were genetically tested and were genotype negative (Sarc -) were used as negative controls and results were normalized to these samples (100% MyBP-C, 100% MYH). Buffer-only controls were tested (0% MyBP-C, 0% MYH). Samples from patients with HCM with a pathogenic truncating *MYBPC3* variant, were utilized as a positive control. These samples have previously been demonstrated to have a reduced MyBP-C/MYH ratio by mass spectrometry.[1] The ratio of normalized MyBP-C/normalized MYH*100% is reported for each sample in triplicate. For quality control, standard curves (200 μg/ml, 2-fold serial dilution, 8 samples in triplicate) were tested for each sample.

**Cardiomyocyte Preparation and Culture**

**Isolation of Neonatal Rat cardiomyocytes (NRVMs):** As previously described [2], the excised ventricles of 1- to 3-day-old Sprague-Dawley rats (Charles River) were processed according to a modified version of the Worthington Neonatal Cardiomyocyte Isolation System (Worthington Biochemical Corporation). Ventricles were minced in ice-cold HBSS and predigested in 1 mg/ml trypsin (Worthington) at 4°C for 6 hours. Tissue was then digested in 30 U/ml purified collagenase (Worthington), dissolved in Media 199 (Invitrogen) with Earle's salts, L-glutamine, 2.2 g/l sodium bicarbonate, 2% penicillin/streptomycin, 25 mM HEPES, and 15% heat-inactivated qualified FBS (Invitrogen), in a Celstir 50-ml jacketed spinner flask (Wheaton) for 45 minutes at 37°C. Digested tissue was triturated and filtered through a 70-μm strainer, and then incubated for 20 minutes at room temperature to further digest partially degraded collagen. Cell suspensions were pre-plated on untreated plastic dishes for 1 hour at 37°C to reduce adherent fibroblast contamination and then filtered through a 40-μm strainer. NRVMs were seeded on plates coated with 5 μg/ml bovine fibronectin (Sigma-Aldrich) in maintenance media (Media 199 with Earle's salts, L-glutamine, 2.2 g/l sodium bicarbonate, 2% penicillin/streptomycin, 25 mM HEPES, and 5% FBS). Medium was changed 24 hours after plating and every subsequent 48 hours unless otherwise noted.

**Induced pluripotent stem cell (iPSC) derived cardiomyocytes (IPSC-CM):** iPSC-CMs used in this study are from a gene-edited iPSC homozygous *MYBPC3* knockout line (*MYBPC3* [–/–]).[3] Stem cell maintenance, cardiomyocyte production, and cell culture were performed as previously described.[3,4] iPSCs were cultured in STEM flex media (Gibco #A3349401) and passaged weekly using EDTA (Invitrogen #15575020) diluted to 0.5 mM in PBS (Gibco#10010- 023). iPSCs were verified to be free of mycoplasma contamination using MycoAlert Detection Kit (Lonza). iPSCs were differentiated into cardiomyocytes using WNT modulation.[5] Stem cell-derived cardiomyocytes (CMs) underwent metabolic selection using glucose-deprived, lactate-containing media for five days as previously described.[6,7] Following lactate purification, CMs were frozen in Cyrostor (Sigma)at a concentration of $2*10^6$ cells/200 μl in liquid nitrogen. Unless otherwise noted, iPSC-CMs were grown on cell culture plates coated with Matrigel (Corning #354277) diluted 1:500 in DMEM/F12 (Gibco, #11330-032) at room temperature for 1 hour. Matrigel/DMEM is removed, and cardiomyocytes are plated using replating media [RPMI+ P/S, 2% FBS, 0.2 μl/mL of 10 mM thiazovivin (TZV, Cayman Chemical, #14245) in sterile DMSO (dimethyl sulfoxide, Sigma #D2650)]. Unless otherwise noted, iPSC-CMs were pooled from 3 independent differentiations. Cells were maintained in RPMI+ P/S [500 ml- RPMI 1640 [+] L-glutamine [+] phenol red, Gibco #11875-093, 5ml of Penn/Strep 10,000 U/ml Gibco #15140122, 10 mL of B27 supplement, serum-free (Life technologies #17504-001].

**Flag-MyBP-C and Turbo-Flag-MyBP-C immunofluorescence (IF):** As previously described[4,8], unpatterned *MYBPC3* [-/-] iPSC-CMs were plated at 150,000 cells per well in 24 well plate (Costar #3526) on PDMS coverslips (Specialty manufacturing, Saginaw, Michigan). 4 days after replating (day 20 of differentiation), flag-*MYBPC3* or turboID-flag-*MYBPC3* pathogenic missense variants were expressed via adenoviral transduction at MOI 10, media was changed 48 hours later. At 96 hours, cells were washed with PBS and fixed for 15 minutes at room temperature in 2% paraformaldehyde (Sigma-Aldrich). Cells were permeabilized in 0.2% Triton X-100 (Sigma-Aldrich) in TBS for 10 minutes. Cells were blocked in 5% normal goat serum (Vector Biolabs) and 1 mg/ml bovine serum albumin (BSA, Sigma) in TBS for 40 minutes at room temperature. All primary, secondary, or labeled streptavidin were prepared in 7.5%

normal goat serum and 1 mg/ml BSA in TBS. Cells were either stained with primary and secondary antibodies (as described below) to visualize flag-MyBP-C localization or with labeled streptavidin (as described below) to visualize the localization of biotinylated proteins. For cells stained with primary and secondary antibodies, a primary antibody solution contained; MyBP-C mouse monoclonal (sc-137180, Santa Cruz) at 1:200, flag rabbit monoclonal (F7425, Millipore) at 1:200 and a secondary antibody solution contained goat anti-mouse IgG Alexa Fluor 488 1:1000 (ThermoFisher Scientific, A11001); and goat anti-rabbit IgG Alexa Fluor 594 1:1000 (ThermoFisher Scientific, A11012). The primary antibody solution was added and incubated overnight at 4 °C, cells were then washed TBS-T for 5 minutes and twice with TBS. Next, the secondary antibody solution was added and incubated for 30 minutes at room air protected from light, and then washed with TBS-T for 5 minutes and twice with TBS. For cells stained with labeled streptavidin, cells were incubated with a solution containing AlexaFlour 488 labeled streptavidin (Invitrogen #S11223) at 1:1000 for 1 hour at room temperature and then washed with TBS-T for 5 minutes and twice with TBS. Next, after primary and secondary antibodies or labeled streptavidin incubations were complete, cells were incubated with 200 ng/ml DAPI in TBS for 10 minutes at room temperature, protected from light. Lastly, coverslips were mounted onto slides with ProLong Diamond Antifade (ThermoFisher Scientific) and cured overnight before imaging. PDMS coverslips were mounted face-up and topped with glass coverslips. Each sample was tested in biologic duplicates and six independent images per coverslip were acquired using a Nikon Eclipse Ti-E inverted fluorescence microscope on 40X Plan Flour DIC M/N2 (Nikon) objective.

**Flag-MyBP-C immunoprecipitation (IP):** Flag-MyBP-C IP was performed as previously described.[2] Briefly, unpatterned neonatal rat ventricular cardiomyocytes (NRVMs) were plated at a density of $1 \times 10^7$ cells per 100-mm culture dish. Cells that were non-transduced (negative control), transduced with WT flag-*MYBPC3* (positive control) or transduced with C3 and C6 pathogenic missense variants flag-*MYBPC3* Arg495Gln, Arg502Trp, Trp792Arg, Arg810His in biologic duplicates using adenovirus (MOI 10) 24 hours after plating. Cells were collected 48 hours later by scraping in ice-cold PBS with Roche protease inhibitor cocktail. Cell pellets were stored at –80°C, and $2 \times 10^7$ cells were used per sample. Pellets were lysed with RIPA buffer (1% Triton X-100, 1% sodium deoxycholate, 0.1% SDS, 150 mM NaCl, 10 mM $Na_2PO_4$ pH 7.2, 1 mM NaF, 1 mM EDTA, 2.5 mM EGTA, 20 mM $(NH_4)_2MoO_4$, 100 μM $Na_3VO_4$) containing protease inhibitor cocktail (complete Mini, Roche) containing phosphatase inhibitor cocktail (PhosSTOP, Roche) and incubated on ice for 15 minutes. Lysates were then sonicated for four 10-second bursts at 50% amplitude using a Branson digital sonifier with cup horn attachment and incubated on ice for another 15 minutes. Protein concentration of whole cell lysate was determined by DC protein assay and diluted to a concentration of 500 μg/100 μl in a total volume of 110 μl. 10 μl of input samples were saved for silver stain analysis, and 100 μl of input samples (500 μg/100 μl) were allocated to 1 FLAG co-IP tube. The second biologic replicate did not have enough total protein to aliquot samples for silver stain analysis. FLAG co-IP tubes received 20 µl anti-FLAG M2–conjugated Sepharose beads (Sigma-Aldrich), washed 3 times in RIPA buffer and were incubated with 500 μg (100 μl) of whole cell lysate. Following overnight incubation at 4°C with gentle shaking, unbound supernatant was collected and beads were washed twice with 100µl of RIPA buffer and once with 200µl of PBS. Flag- immunoprecipitated sample was eluted by competitive binding in 100µl with 100µg/ml 3X FLAG peptide (Sigma-Aldrich) for 2 hours. Ninety µl of the elution sample was submitted to mass spectrometry and 10 µl was saved for western blot analysis.

**Flag Western Blot and Silver Stain Analysis of Flag-IP:** For gel analysis, the immunoprecipitation samples from the first biologic replicate were analyzed. 10 μl of cellular lysate (input) and 10 μl flag immunoprecipitated (flag-IP elution) samples were each mixed with 3 μl of 4x Laemmli sample buffer (Biorad) and 1μl of -Mercaptoethanol (Sigma Aldrich) and heated to 95°C for 5 minutes. 5 μl/well of input samples was loaded on Criterion Precast gel 4-20% (BioRad 3450034) and run with Running buffer (6.06 g Tris Base, 28.2 g Glycine, 2 g SDS) at 60 mA for 1 hour. Silver stain (Sigma-PROTSIL2) was performed using manufacturer's protocol. 5 μl/well of elution samples were loaded on Criterion Precast gel 4-20% (BioRad 3450034) and run with Running buffer (6.06 g Tris Base, 28.2 g Glycine, 2 g SDS) at 60 mA for 1 hour. After performing a wet transfer to nitrocellulose (Biorad) a western blot was performed using LICOR blocking buffer to block the membrane for 1 hour. The nitrocellulose membranes was then incubated with LICOR and antibody diluent buffer with the primary antibody anti-flag, mouse monoclonal (F1804 Millipore) at 1:200 incubated overnight at 4$^{o}$C, the membrane was then washed three times for 5 minutes with PBS-T and incubated with a secondary antibody goat anti-mouse IRDye800CW (Licor, 926-3222100) at 1:10,000 protected from light for 1 hour, and then washed three times again for 5 minutes with PBS-T before imaging. The membrane was imaged using Odyssey CLx Licor.

**Protein identification of flag-IP samples using LC-MS/MS:** LC-MS/S of flag-IP samples were performed by MS Bioworks (Ann Arbor, MI). Thirty microliters of the flag-IP (elutant samples) were separated by 10% Bis-Tris Novex mini-gel (Invitrogen) using the MES buffer system. The gel was stained with Coomassie, and each lane was excised into ten equally sized segments. Gel pieces were processed using a robot (ProGest, DigiLab) with the following protocol: Washed with 25mM ammonium bicarbonate followed by acetonitrile, reduced with 10mM dithiothreitol at 60°C followed by alkylation with 50mM iodoacetamide at RT, digested with trypsin (Promega) at 37°C for 4h, quenched with formic acid, and the supernatant was analyzed directly without further processing.

**Data-dependent acquisition (DDA) method:** The gel digests were analyzed by nano LC/MS/MS with a Waters M-class HPLC system interfaced to a ThermoFisher Fusion Lumos. Peptides were loaded on a trapping column and eluted over a 75µm analytical column at 350nL/min; both columns were packed with Luna C18 resin (Phenomenex). A 30min gradient was employed. The mass spectrometer was operated in data-dependent mode, with MS and MS/MS performed in the Orbitrap at 60,000 FWHM resolution and 15,000 FWHM resolution, respectively. Advanced peak determination was turned on. The instrument was run with a 3s cycle for MS and MS/MS. Data was collected for 2 independent co-IP-MS experiments. Data were searched using a local copy of Mascot with the following parameters: Enzyme: Trypsin, Database: Uniprot Rat (forward and reverse appended with common contaminants and MYBPC3_HUMAN), Fixed modification: Carbamidomethyl (C), Variable modifications: Oxidation (M), Acetyl (Protein N-term), Deamidation (NQ), Pyro-Glu (N-term Q), Mass values: Monoisotopic Peptide Mass Tolerance: 10 ppm, Fragment Mass Tolerance: 0.02 Da, Max Missed Cleavages: 2. Mascot DAT files were parsed into the Scaffold software for validation, filtering and to create a nonredundant list per sample. Data were filtered 1% protein and peptide level false discovery rate (FDR) and required at least two unique peptides per protein. The full list of proteins was evaluated, and known contaminants were excluded, resulting in unnormalized spectral counts. These were then converted to normalized spectral abundance factors (NSAF). Based on the equation.

$NSAF = (SpC/MW)/\Sigma(SpC/MW)_N$
Where SpC = Spectral Counts
MW = Protein MW in kDa
N = Total Number of Proteins

**Defining MYBPC3 interacting proteins by LC-MS/MS**: Criteria for identifying *MYBPC3* interacting proteins were the following: (a) proteins were detected in both experimental replicates for a given sample, (b) proteins showed a calculated fold-change score in normalized spectral abundance factor (NSAF) of 2 or higher over the non-transduced samples.

**Identifying changes in MyBP-C interacting proteins in the presence of pathogenic missense variants:** Criteria for proteins differentially associated with MyBP-C in the presence of pathogenic missense variants is as follows (a) proteins showed a 50% increase or decrease in NSAF compared to WT MyBP-C samples, and exhibited (b) FDR adjusted p-value (q-value) < 0.05, p-values were calculated using a two-tailed 2 -sample t-test and were corrected for multiple comparisons by calculating q-value using a two-stage step up method of Benjamini, Krieger, and Yekutieli. We limited our analysis to proteins identified as *MYBPC3* interacting proteins. NSAF were compared for WT samples and individual pathogenic variants. We also compared all pathogenic variants as a single group (WT vs pathogenic variants- Arg495Gln, Arg502Trp, Trp792Arg, Arg810His) and pathogenic variants within a given subdomain (WT vs C3 pathogenic variants- Arg495Gln and Arg502Trp samples, WT vs C6 pathogenic variants- Trp792Arg and Arg810His variants).

**Biotin-ligase proximity labeling (PL):**The protocol for Biotin-ligase PL of WT turboID-flag-MyBP-C and four pathogenic variants (Arg495Gln, Arg502Trp, Trp792Arg, Arg810His) is as follows. In a 6-well tissue culture plate (Fisher, FB012927), wells were coated with 0.75 mL of bovine fibronectin (Sigma, 1mg/mL) diluted 1:200 in sterile HBSS (Corning) for 1 hour. Unpatterned iPSC-CMs gene-edited to be a homozygous *MYBPC3* knockout line (*MYBPC3* [–/–])[3] were plated at a cellular density of 2 x $10^6$ cells per well in replating media (as described in supplemental materials). Cells were cultured until day 20 of differentiation, maintained in 2ml/well of RPMI + P/S, with media changes occurring every 48 hours. On day 20 of differentiation iPSC-CMs were treated with adenovirus at MOI 10 for 48 hours, carrying one of the following constructs: flag-MYBPC3 – negative control, turboID-flag-MYBPC3 WT – positive control, or a turboID-flag MYBPC3 pathogenic mutant: Arg495Gln, Arg502Trp Trp792Arg, or Arg810His. Each sample was tested in 3 biological replicates. At 48 hours, a media change was performed, and on day 4 post-infection (day 24 of differentiation), biotin labeling was performed by changing the media to RPMI + P/S with 50 μM biotin (Sigma B4501) and 500 μM ATP (Sigma) prepared freshly. To prepare biotin labeling media, stocks of Biotin at 20 mM are utilized that contain 100 mg biotin and 2 ml of 30% NH4OH dissolved on ICE and then 5 mL of 1N HCl is slowly added until a total concentration of 18 mL is reached and biotin is dissolved. Biotin can easily precipitate if pH is raised too quickly, so keeping the sample adequately chilled is important. These aliquots are stored protected from light at 4°C for up to 6 months. Buffers are filtered using 0.22 μm PES filter (Millipore)and filter-tipped and ultrapure HPLC grade water (Alfa Aesar) is utilized for all buffers and reagents. After 1 hour of biotin labeling, cells were washed 3 times with PBS (Gibco) and lysed by adding 500 μl/well of lysis buffer (50 mM Tris (Sigma), PH 7.5, 150 mM NaCl (Sigma), 0.4% SDS (Sigma), 1% NP-40 (Thermo), 1.5 mM $MgCl_2$ (Sigma), 1mM EGTA (Sigma), 0.5 mM EDTA (Sigma), protease inhibitor (complete Mini protease inhibitor cocktail, Roche), phosphatase inhibitor cocktail (PhosSTOP, Roche), 200μM Phenylmethylsulfonyl fluoride (PMSF, Thermo), and 500 U benzonase/ml (EMD MIllipore) and cells are scrapped. The collected total cell lysate was sonicated for four 10-second bursts at 50% amplitude using a Branson digital sonifier with cup horn attachment and incubated on ice for a further 15 minutes. Ninety μl of lysate was flash frozen in liquid nitrogen and stored at- 80°C for western blot analysis and 10 μl of lysate were utilized for protein concentration determination using DC protein assay.

The protein concentration was determined for the remaining 400 μl of lysate and 100 μg of protein per sample was added to 750 μl of dynabeads myOne streptavidin C1 beads (Invitrogen), and prewashed three times with lysis buffer within Lobind 1.5 ml microcentrifuge tubes. Lysate was incubated with streptavidin beads on a rotator (end over end) overnight at 4°C. Each sample's beads were washed with 1ml lysis buffer once, 1ml wash buffer (50 mM Tris, pH 7.5) once, 1ml lysis buffer twice, and 50 mM ammonium bicarbonate (Sigma, pH 8.0 in water) six times. Beads were separated from supernatants using dynamag-2 (magnetic holder for Eppendorf tubes, Fisher 12321D) and all samples were saved. Beads are then resuspended in 1 mL of ammonium bicarbonate (pH 8.0 in water) and 100 μl was removed for western blot analysis (elutant). For the remaining 900 μl of beads utilized for mass spectrometry analysis, ammonium bicarbonate was removed and the beads were stored at -80°C.

**Western Blot and Silver Stain Analysis of MyBP-C PL:** 10 μl of cellular lysate (input) was mixed with 3 μl of 4x Laemmli sample buffer (biorad) and 1μl of 2-Mercaptoethanol (BME, Sigma Aldrich) and heated to 95°C for 5 minutes. Streptavidin beads (elutant) samples saved for gel analysis were resuspended in 100 ul of 1x Laemmli sample buffer (biorad) with 200 μM biotin and 5 μl of BME and heated to 95°C for 5 minutes. Supernatant was separated from beads using dynamag-2. 5 μl/well of input samples and 10 μl/well of elutant samples were loaded on Citerion Precast gel 4-20% (BioRad 3450034) and run with Running buffer (6.06 g Tris Base, 28.2 g Glycine, 2 g SDS) at 60 mA for 1 hour. After performing a wet transfer to nitrocellulose (Biorad) a western blot was performed using LICOR blocking buffer to block the membrane for 1 hour. The nitrocellulose membranes were then incubated with Streptavidin IRDye 800 antibody (LICOR) at 1:1,000 for 1 hour (protected from light) and then washed three times for 5 minutes with PBS-T. Silver stain (Sigma-PROTSIL2) was performed using manufacturer's protocol. Gels were imaged using Odyssey CLx Licor.

**Protein identification of MyBP-C PL samples using LC-MS/MS:** LC-MS/MS was performed by MS Bioworks (Ann Arbor, MI). On-bead trypsin digestion was performed manually with the following protocol: The beads were washed x2 with 25mM ammonium bicarbonate. Proteins were reduced by 10mM dithiothreitol at 60°C followed by alkylation with 15mM iodoacetamide at RT. Supernatant was removed and proteins were digested with 100ng (10μL x 10ng/μL) trypsin (Promega) at 37°C for 18h. A further 100 ng trypsin was added and incubated for 4h. Digestion was quenched with formic acid, and the supernatant was desalted. Peptides were taken to dryness using a lyophilizer. Peptides were reconstituted in 0.1% TFA for mass spectrometry analysis. **T**en percent of each sample was analyzed in analytical triplicate by LC/MS with a ThermoFisher Vanquish Neo UPLC system interfaced to a ThermoFisher Orbitrap Astral using DDA, and another ten percent of each sample using DIA. Peptides were loaded on a trapping column and eluted over a 75μm analytical column at 350nL/min; the column was heated to 55 °C. A 30 min gradient was employed**.**

**Identifying MyBP-C proximity proteins using LC-MS/MS data-dependent acquisition (DDA):** We first evaluated if proteins were proximity proteins with MyBP-C (WT or pathogenic variants- Arg495Gln, Arg502Trp, Trp792arg, Arg810His). The mass spectrometer was operated in data-dependent acquisition (DDA) mode, with MS performed in the Orbitrap at 120,000 FWHM resolution and MS/MS performed in the Astral at 80,000 FWHM resolution, respectively. Advanced Peak Determination was turned on. The instrument was run with a top 200 method. Data was processed using the Proteome Discover 3.1 which served several functions: Recalibration of MS data, filtering of database search results at the 1% protein and peptide false discovery rate (FDR), calculation of MS1 peak areas and PSMs per sample. Data were search using Chimerys with the following parameters: Enzyme: Trypsin, Database: Swissprot Human, fixed modification: Carbamidomethyl (C), Variable modifications: Oxidation (M), Fragment Mass tolerance 200 ppm. The result file was uploaded to perseus v1.5.5.3 for further analysis. The full list of proteins was reported as unnormalized spectral counts (SpC). These were then converted to normalized spectral abundance factors (NSAF). Based on the equation.

$NSAF = (SpC/MW)/\Sigma(SpC/MW)_N$
Where SpC = Spectral Counts
MW = Protein MW in kDa
N = Total Number of Proteins

**Defining MyBP-C proximity proteins by LC-MS/MS**

Criteria for identifying *MYBPC3* proximity proteins were the following: (a) proteins were detected in all experimental and technical replicates, and (b) proteins showed a calculated fold-change score in normalized spectral abundance factor (NSAF) of 2 or higher over the negative control samples or were absent in the negative controls. To create volcano plots, data were processed using log2 transforming intensity values.

**Changes in MyBP-C proximity, using LC-MS/MS data-independent acquisition (DIA):** We utilized data-independent acquisition (DIA) mass spectrometry to identify changes in MyBP-C proximity proteins between WT MyBP-C and individual MyBP-C carrying pathogenic. The mass spectrometer was operated in data-independent mode. Sequentially, full scan MS data (240,000 FWHM resolution) from m/z 380-980 was followed by 300 x 2m/z precursor isolation windows; products were acquired in the Astral at 80,000 FWHM resolution. The maximum ion inject time (IIT) was set to 3.5ms for DIA; the NCE was set to 25. A pool was created from 10% of all three samples. An injection of the pool was included at the start and end. DIA data were analyzed suing DIA-NN (v.1.9.1) which severed several functions: conversion of RAW files to QUANT, alignment based on retention times, searching data using *in-silico* spectral library created from the FASTA file and iteratively using the spectral library created from RAW data, filtering of database search results at 1% peptide and protein false discovery rate (FDR), calculation of peak areas for detected peptides, data normalization. Data were searched using Chimerys with the following parameters: Enzyme Trypsin, Data Base: Swissprot Human, Fixed modification: Carbamidomethyl (C), Variable modification: oxidation (M), precursor mass tolerance: determined, Fragment mass tolerance: determined, Peptide FDR 0.01 (1%), Protein FDR 0.01 (1%), peptide length 7-30 AA, max missed cleavages: 1, min peptides: 1 The report.pg_matrix.tsv output was uploaded to Perseus v1.5.5.3 for further analysis. Proteins were reported as normalized protein intensity. Data was processed log2 transforming intensity values with missing values replaced with 0.

**Identifying proteins with altered MyBP-C proximity in the presence of pathogenic missense variants:** Differences in normalized intensity values were compared for proteins present in all WT and pathogenic variant samples and were identified by performing a student's 2-tailed 2-sample t-test. P-values were corrected for multiple comparisons by calculating q-value with Benjamini-Hochberg FDR correction (FDR 5%). Proteins with altered proximity were defined as proteins with a fold change of log2 transformed values > 0.5 or < -0.5 (increase or decrease in *MYBPC3* binding) and FDR adjusted p-value (q-value) < 0.05.

**Gene Ontology Enrichment Analysis:** Gene ontology analysis was performed using metascape [http://metascape.org].[9] We analyzed the following lists (1) interacting/proximity proteins (2) proteins with increased association/proximity to pathogenic variant(s) (3) proteins with decreased association/proximity to pathogenic variant(s). For the flag- immunoprecipitation mass spectrometry which was performed in neonatal rat ventricular cardiomyocytes we used the rattus norvegicus reference and queried the following annotated data sets (1) GO biological process complete (2) GO molecular function complete (3) GO cellular component complete (4) Reactome pathway (Reactome version 86 Released 2023-09-07). Pathway and process enrichment analyses were performed as follows:

We first identified all statistically enriched terms (can be GO terms, reactomepathways), accumulative hypergeometric p-values and enrichment factors were calculated and used for filtering. Remaining significant terms were then hierarchically clustered into a tree based on Kappa-statistical similarities among their gene memberships (similar to what is used in NCI DAVID site). We used a min overlap 3, p-value cutoff < 0.01, and min enrichment 1.5. Then 0.3 kappa score was applied as the threshold to cast the tree into term clusters. The terms within each cluster are exported in the Excel spreadsheet named "Enrichment Analysis". Results of this analysis were presented as follows; The Circos plot shows how genes from the input gene lists overlap. We selected the term with the best p-value within each cluster as its representative term and display them in a dendrogram. The dendrogram cells are colored by their -log 10 (p-values), grey cells indicate the lack of enrichment for that term in the corresponding gene list. Protein-protein interactions among input genes were extracted by Metascape from PPI data source and formed a PPI network. MCODE algorithm was then applied to this network to identify neighborhoods where proteins are densely connected. GO enrichment analysis was applied to each MCODE network to extract "biological meanings" from the network component, where top three best p-value terms were retained. For proximity-labeling DDA mass spectrometry on WT MyBP-C proximity proteins GO enrichment analysis was also performed using a second GO enrichement software (TOPPGENE)[10] for cellular compartment and molecular function. Gene categories with a Bonferroni corrected p-value of 0.05 or lower were included.

Supplemental Table 1: MYBPC3 non-synonymous missense variants identified in patients with HCM who underwent genetic testing within the ShaRe registry (2025Q2).

File: SupplementalTable1.xlsx

Supplemental Table 2: AQUA Peptides

| AQUA peptides | |
|---|---|
| R502W (Elastase) | R810H/W792R (Trypsin) |
| REETFKYWF**K** (heavy)<br>-W502 mutant | ISNVGEDSC[+57.021464]TVQWEPPAYDGGQPILGYILEH**K** (heavy)<br>- H810 mutant |
| REETFKYWFK**K**<br>(heavy)<br>-W502 mutant | ISNVGEDSC[+57.021464]TVQWEPPAYDGGQPILGYILE**R** (heavy)<br>- WT |
| REETFKY**R** (heavy)- -<br>R502 WT | ISNVGEDSC[+57.021464]TVQ**R** (heavy)<br>- R792 mutant |
| | G531R (Trypsin) |
| | HHLIINEAMLEDAGHYAL**C**TS**R** (heavy) -R531 Mutant |
| | HHLIINEA**M**LEDAGHYAL**C**TS**R** (heavy) – R531 Mutant |
| | HHLIINEAMLEDAGHYAL**C**TSGGQALAELIVQE**K** (heavy) – G531 WT |
| | HHLIINEA**M**LEDAGHYAL**C**TSGGQALAELIVQE**K** (heavy) – G531 WT |

Supplemental Table 3: Peptide m/z values used in AQUA mass spectrometry

| R502W (Elastase) | | | R810H/W792R(trypsin) | | |
|---|---|---|---|---|---|
| Compound | m/z | z | Compound | m/z | z |
| REETFKYWFK (light) | 478.579 | 3 | SNVGEDSC[+57.021464]TVQWEPPAYDGGQPILGYILEHK (light) | 1191.572 | 3 |
| REETFKYWFK (heavy) | 481.25 | 3 | ISNVGEDSC[+57.021464]TVQWEPPAYDGGQPILGYILEHK (heavy) | 1194.243 | 3 |
| REETFKYWFKK (light) | 521.277 | 3 | ISNVGEDSC[+57.021464]TVQWEPPAYDGGQPILGYILER (light) | 1155.221 | 3 |
| REETFKYWFKK (heavy) | 523.948 | 3 | ISNVGEDSC[+57.021464]TVQWEPPAYDGGQPILGYILER (heavy) | 1158.557 | 3 |
| REETFKYR (light) | 376.865 | 3 | ISNVGEDSC[+57.021464]TVQR (light) | 732.841 | 2 |
| REETFKYR (heavy) | 380.201 | 3 | ISNVGEDSC[+57.021464]TVQR (heavy | 737.8451 | 2 |
| | | | G531R (trypsin) | | |
| | | | HHLIINEAMLEDAGHYALC[+57.021464]TSR (light) | 638.56 | 4 |
| | | | HHLIINEAMLEDAGHYALC[+57.021464]TSR (heavy) | 641.062 | 4 |
| | | | HHLIINEAM[+15.994915]LEDAGHYALC[+57.021464]TSR (light) | 642.5587 | 4 |
| | | | HHLIINEAM[+15.994915]LEDAGHYALC[+57.021464]TSR (heavy) | 645.0608 | 4 |
| | | | HHLIINEAMLEDAGHYALC[+57.021464]TSGGQALAELIVQEK (light) | 933.7185 | 4 |
| | | | HHLIINEAMLEDAGHYALC[+57.021464]TSGGQALAELIVQEK (heavy) | 935.722 | 4 |
| | | | HHLIINEAM[+15.994915]LEDAGHYALC[+57.021464]TSGGQALAELIVQEK (light) | 937.7172 | 4 |
| | | | HHLIINEAM[+15.994915]LEDAGHYALC[+57.021464]TSGGQALAELIVQEK (heavy) | 939.7207 | |

Supplemental Table 4: Flag-immunoprecipitation MyBP-C interacting proteins

File: SupplementalTable4_FlagIP_IP.xlsx

Supplemental Table 5: Differentially Associated (DA) Flag-MyBP-C interacting proteins individual variants

File: SupplementalTable5_DA_IP.xlsx

Supplemental Table 6: Differentially Associated (DA) Flag-MyBP-C, comparing grouped pathogenic variants to WT MyBP-C

File: SupplementalTable6_DA_groupIP.xlsx

**Supplemental Table 7: Proximity labeling mass spectrometry DDA overview of protein totals**

| Sample | Technical Replicate 1 # | Technical Replicate 1 Int | Techincal Replicate 2 # | Techincal Replicate 2 Int | Technical Replicate 3 # | Technical Replicate 3 Int | Average # | Average Int | Standard deviation # | Standard deviation Int |
|---|---|---|---|---|---|---|---|---|---|---|
| Neg-1 | 1366 | 3.20E+04 | 1399 | 2.71E+04 | 1478 | 2.75E+04 | 1414 | 2.89E+04 | 58 | 3.E+03 |
| Neg-2 | 1210 | 2.23E+04 | 665 | 1.05E+04 | 623 | 1.05E+04 | 833 | 1.44E+04 | 327 | 7.E+03 |
| Neg-3 | 928 | 1.89E+04 | 1001 | 2.31E+04 | 1050 | 2.20E+04 | 993 | 2.13E+04 | 61 | 2.E+03 |
| **WT-1** | **3997** | **1.66E+05** | **4003** | **1.69E+05** | **4012** | **1.71E+05** | **4004** | **1.69E+05** | **8** | **3.E+03** |
| **WT-2** | **3762** | **1.07E+05** | **3784** | **1.06E+05** | **3806** | **1.08E+05** | **3784** | **1.07E+05** | **22** | **6.E+02** |
| **WT-3** | **3098** | **8.56E+04** | **3083** | **7.85E+04** | **3088** | **8.41E+04** | **3090** | **8.27E+04** | **8** | **4.E+03** |
| R495Q-1 | 2290 | 2.38E+04 | 2241 | 2.40E+04 | 2371 | 2.38E+04 | 2301 | 2.39E+04 | 66 | 1.E+02 |
| R495Q-2 | 2922 | 5.76E+04 | 2816 | 5.50E+04 | 2909 | 5.46E+04 | 2882 | 5.57E+04 | 58 | 2.E+03 |
| R495Q-3 | 3960 | 1.56E+05 | 3985 | 1.56E+05 | 3974 | 1.60E+05 | 3973 | 1.57E+05 | 13 | 2.E+03 |
| R502W-1 | 3931 | 1.48E+05 | 3980 | 1.49E+05 | 3975 | 1.49E+05 | 3962 | 1.49E+05 | 27 | 2.E+02 |
| R502W-2 | 4048 | 1.86E+05 | 4050 | 1.89E+05 | 4049 | 1.92E+05 | 4049 | 1.89E+05 | 1 | 3.E+03 |
| R502W-3 | 4019 | 1.80E+04 | 4028 | 1.80E+05 | 3996 | 1.79E+05 | 4014 | 1.26E+05 | 17 | 9.E+04 |
| W792R-1 | 4039 | 1.93E+05 | 4039 | 1.97E+05 | 4048 | 1.97E+05 | 4042 | 1.96E+05 | 5 | 2.E+03 |
| W792R-2 | 4039 | 1.74E+05 | 4045 | 1.73E+05 | 4041 | 1.76E+05 | 4042 | 1.74E+05 | 3 | 2.E+03 |
| W792R-3 | 4043 | 1.83E+05 | 4061 | 1.86E+05 | 4033 | 1.83E+05 | 4046 | 1.84E+05 | 14 | 2.E+03 |
| R810H-1 | 4039 | 1.90E+05 | 4009 | 1.95E+05 | 4027 | 1.91E+05 | 4025 | 1.92E+05 | 15 | 3.E+03 |
| R810H-2 | 4018 | 1.80E+05 | 4024 | 1.81E+05 | 4019 | 1.83E+05 | 4020 | 1.81E+05 | 3 | 1.E+03 |
| R810H-3 | 423 | 2.84E+03 | 392 | 2.37E+03 | 380 | 2.34E+03 | 398 | 2.51E+03 | 22 | 3.E+02 |

The number of proteins (#) and intensity (Int) are reported for each biological replicate tested in technical triplicates .One biologic replicate of Arg810His (indicated in red) had a low level of protein detection in both DIA and DDA mass spectrometry, comparable to the negative control (flag-MyBP-C), suggesting failure of on-bead trypsinization. This sample was excluded from further analysis.

**Supplemental Table 8: Proximity labeling mass spectrometry DIA overview of protein totals**

| Sample | Technical Replicate 1 | | Technical Replicate 2 | | Technical Replicate 3 | | Average | | Standard deviation | |
|---|---|---|---|---|---|---|---|---|---|---|
| | # | Int | # | Int | # | Int | # | Int | # | Int |
| Neg-1 | 1965 | 3.20E+10 | 2019 | 5.10E+10 | 2142 | 4.80E+10 | 2042 | 4.37E+10 | 2068 | 4.76E+10 |
| Neg-2 | 1074 | 2.90E+10 | 1016 | 3.50E+10 | 970 | 3.50E+10 | 1020 | 3.30E+10 | 1002 | 3.43E+10 |
| Neg-3 | 1694 | 2.40E+10 | 1801 | 3.30E+10 | 1678 | 2.60E+10 | 1724 | 2.77E+10 | 1734 | 2.89E+10 |
| **WT-1** | **7375** | **2.80E+10** | **7467** | **2.80E+10** | **7459** | **2.70E+10** | **4950** | **2.77E+10** | **4141** | **2.76E+10** |
| **WT-2** | **6941** | **2.60E+10** | **6948** | **2.60E+10** | **6957** | **2.60E+10** | **6949** | **2.60E+10** | **6951** | **2.60E+10** |
| **WT-3** | **5525** | **2.50E+10** | **5471** | **2.60E+10** | **5447** | **2.60E+10** | **5481** | **2.57E+10** | **5466** | **2.59E+10** |
| R495Q-1 | 3973 | 2.30E+10 | 3893 | 2.60E+10 | 3841 | 2.50E+10 | 3902 | 2.47E+10 | 3879 | 2.52E+10 |
| R495Q-2 | 4828 | 2.50E+10 | 4684 | 2.60E+10 | 4816 | 2.70E+10 | 4776 | 2.60E+10 | 4759 | 2.63E+10 |
| R495Q-3 | 7231 | 2.90E+10 | 7266 | 2.90E+10 | 7270 | 2.80E+10 | 7256 | 2.87E+10 | 7264 | 2.86E+10 |
| R502W-1 | 7328 | 2.80E+10 | 7403 | 2.70E+10 | 7390 | 2.70E+10 | 7374 | 2.73E+10 | 7389 | 2.71E+10 |
| R502W-2 | 7604 | 2.80E+10 | 7641 | 2.80E+10 | 7646 | 2.70E+10 | 7630 | 2.77E+10 | 7639 | 2.76E+10 |
| R502W-3 | 7499 | 2.80E+10 | 7494 | 2.80E+10 | 7494 | 2.80E+10 | 7496 | 2.80E+10 | 7495 | 2.80E+10 |
| W792R-1 | 7605 | 2.80E+10 | 7587 | 2.80E+10 | 7588 | 2.80E+10 | 7593 | 2.80E+10 | 7589 | 2.80E+10 |
| W792R-2 | 7580 | 2.80E+10 | 7591 | 2.70E+10 | 7586 | 2.70E+10 | 7586 | 2.73E+10 | 7588 | 2.71E+10 |
| W792R-3 | 7510 | 2.80E+10 | 7476 | 2.80E+10 | 7518 | 2.80E+10 | 7501 | 2.80E+10 | 7498 | 2.80E+10 |
| R810H-1 | 7558 | 2.90E+10 | 7571 | 2.90E+10 | 7577 | 2.80E+10 | 7569 | 2.87E+10 | 7572 | 2.86E+10 |
| R810H-2 | 7481 | 2.90E+10 | 7481 | 2.80E+10 | 7450 | 2.90E+10 | 7471 | 2.87E+10 | 7467 | 2.86E+10 |
| R810H-3 | 922 | 7.80E+10 | 766 | 1.40E+11 | 516 | 1.40E+11 | 735 | 1.19E+11 | 672 | 1.33E+11 |

The number of proteins (#) and intensity (Int) are reported for each biological replicate tested in technical triplicates. One biologic replicate of Arg810His (indicated in red) had a low level of protein detection in both DIA and DDA mass spectrometry, comparable to the negative control (flag-MyBP-C), suggesting failure of on-bead trypsinization. This sample was excluded from further analysis.

Supplemental Table 9: TurboID proximity labeling and DDA mass spectrometry identifies MyBP-C proximity proteins.

File: SupplementalTable9_PP_DDA.xlsx

Supplemental Table 10: comparison of Flag-IP MyBP-C interacting proteins (IP, 252) and Proximity labeling MyBP-C proximity proteins (PP 3,240) results DDA mass spectrometry

File SupplementalTable10_comparison_IP_PP_DDA.xlsx

Supplemental Table 11: TurboID proximity labeling and DIA mass spectrometry identifies MyBP-C proximity proteins with decreased or increased proximity to MyBP-C in the presence of P/LP *MYBPC3* missense variant.

File: SupplementalTable11_DP_DIA.xlsx

Supplemental Table 12: Comparison of differential MyBP-C association and differential MyBP-C proximity

File: SupplementalTable12_comparison_DA_DP.xlsx

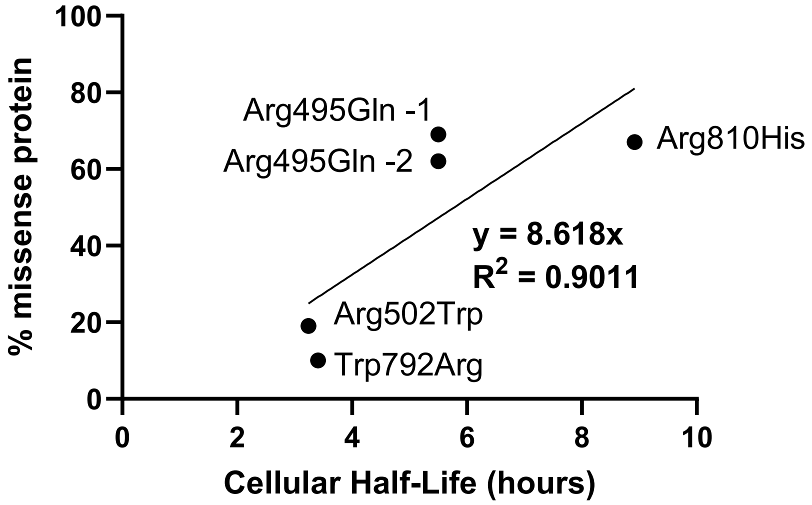


**Supplemental Figure 1: Linear correlation between previously measured cellular half-lives and % mutant protein measured by AQUA in human left ventricular tissue.** Correlation between mutant protein fraction from current AQUA data (Figure 1C) combined with prior AQUA data on two mutant protein fraction for two Arg495Gln patient samples (Y axis) [11] and previously measured cellular lives [8]. Linear regression was performed, assuming y-intercept of 0.

A)

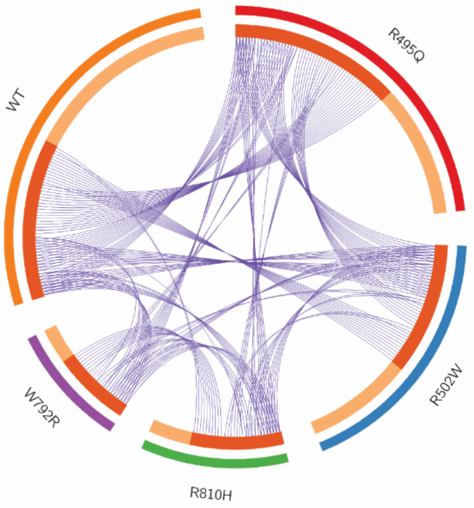


B)

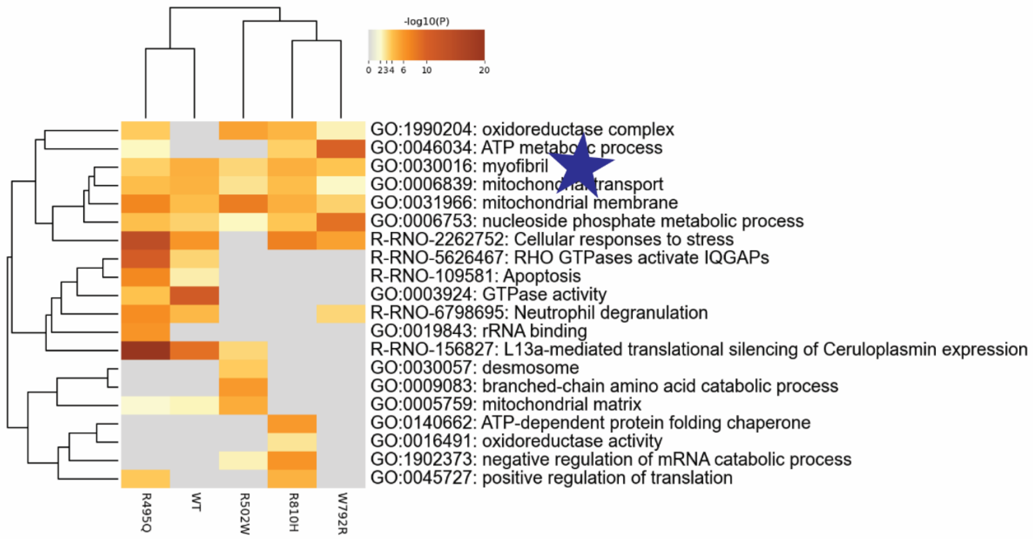


**Supplemental Figure 2: Flag-immunoprecipitation MyBP-C interacting proteins.** A) 252 interacting proteins were identified: 116 for MyBP-C WT, 98 for MyBP-C Arg495Gln, 79 MyBP-C for Arg502Trp, and 37 MyBP-C Trp792Arg, and 43 MyBP-C for Arg810His. The Circos plot shows how genes from the input gene lists of interacting proteins overlap. On the outside, each arc represents the identity of each gene list (WT- orange, Arg495Gln-red, Arg502Trp-blue, Arg810His-green, Trp792Arg-purple). On the inside, each arc represents a gene list, where each gene member of that list is assigned a spot on the arc. Dark orange color represents the genes that are shared by multiple lists and light orange color represents genes that are unique to that gene list. Purple lines link the same gene that are shared by multiple gene lists. B) **Gene ontology enrichment analysis of MyBP-C interacting proteins.** Using metascape, we identified all statistically enriched terms and selected the term with the best p-value within each cluster as its representative term and displayed them in a dendrogram. The dendrogram cells are colored by their p-values; grey cells indicate the lack of enrichment for that term in the corresponding gene list. Gene ontology analysis reveals that there is also overlap in the pathways enriched among MyBP-C interacting proteins for both wild-type and pathogenic missense samples, with all samples showing enrichment in myofibril proteins (GO:0030016).

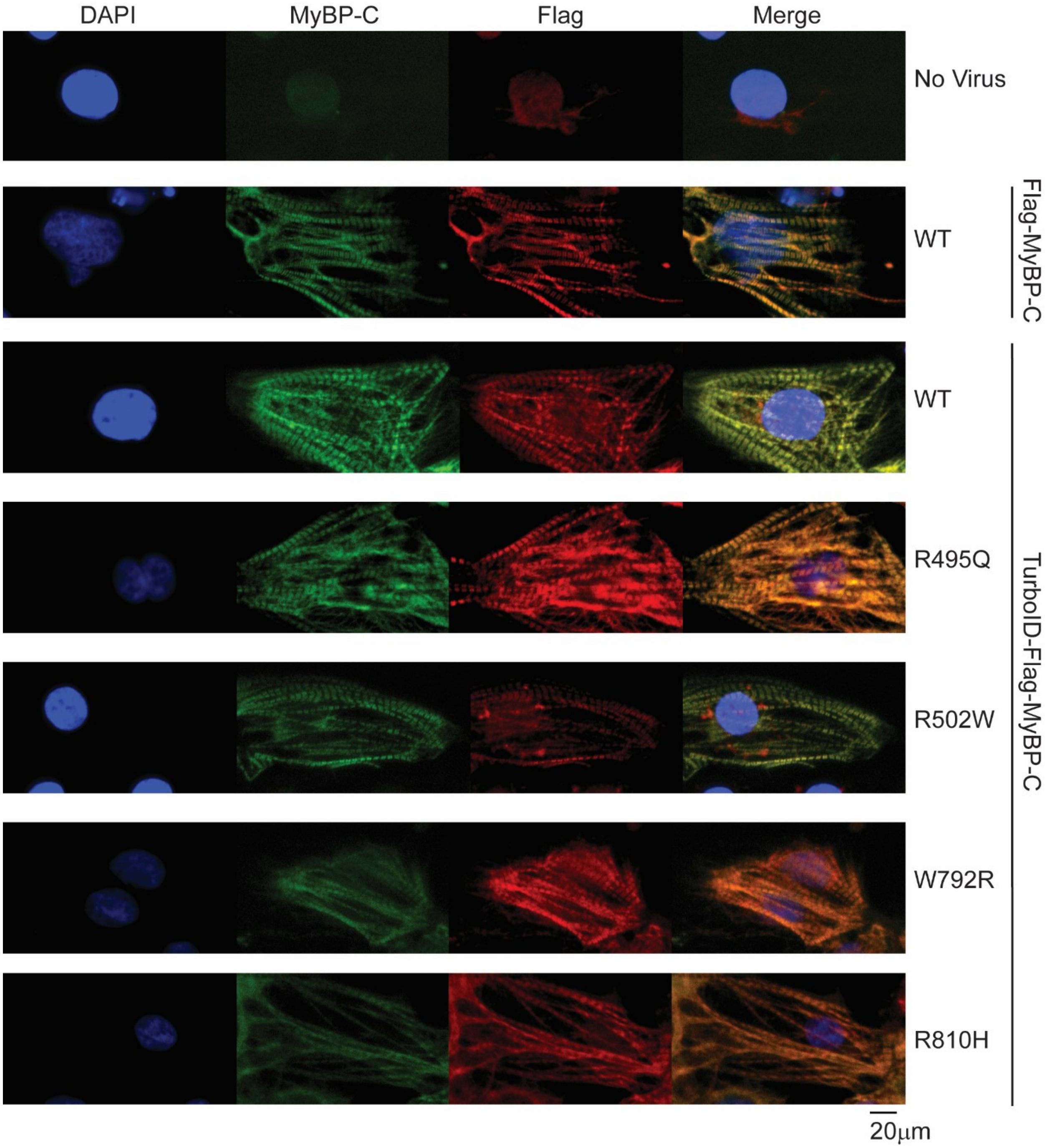


**Supplemental Figure 3: Immunofluorescence of MyBP-C protein constructs in MYBPC3 [-/-] iPSC-CMs.** Localization of various MyBP-C protein constructs [flag-MyBP-C WT and Turbo-ID-flag-MyBP-C WT, Arg495Gln, Arg502Trp, Trp792Arg, and Arg810His], expressed using adenoviral transduction, were evaluated in MYBPC3 [-/-] IPSC-CMs. Non-transduced MYBPC3 [-/-] IPSC-CMs were tested as a negative control. Cells underwent DAPI staining (blue) immunofluorescence using a MyBP-C (green) and flag (red) antibody. Both MyBP-C and Flag immunofluorescence demonstrated that all constructs localize normally within myofibrils. Samples were tested in biological duplicates, and 6 representative images were collected from each sample.

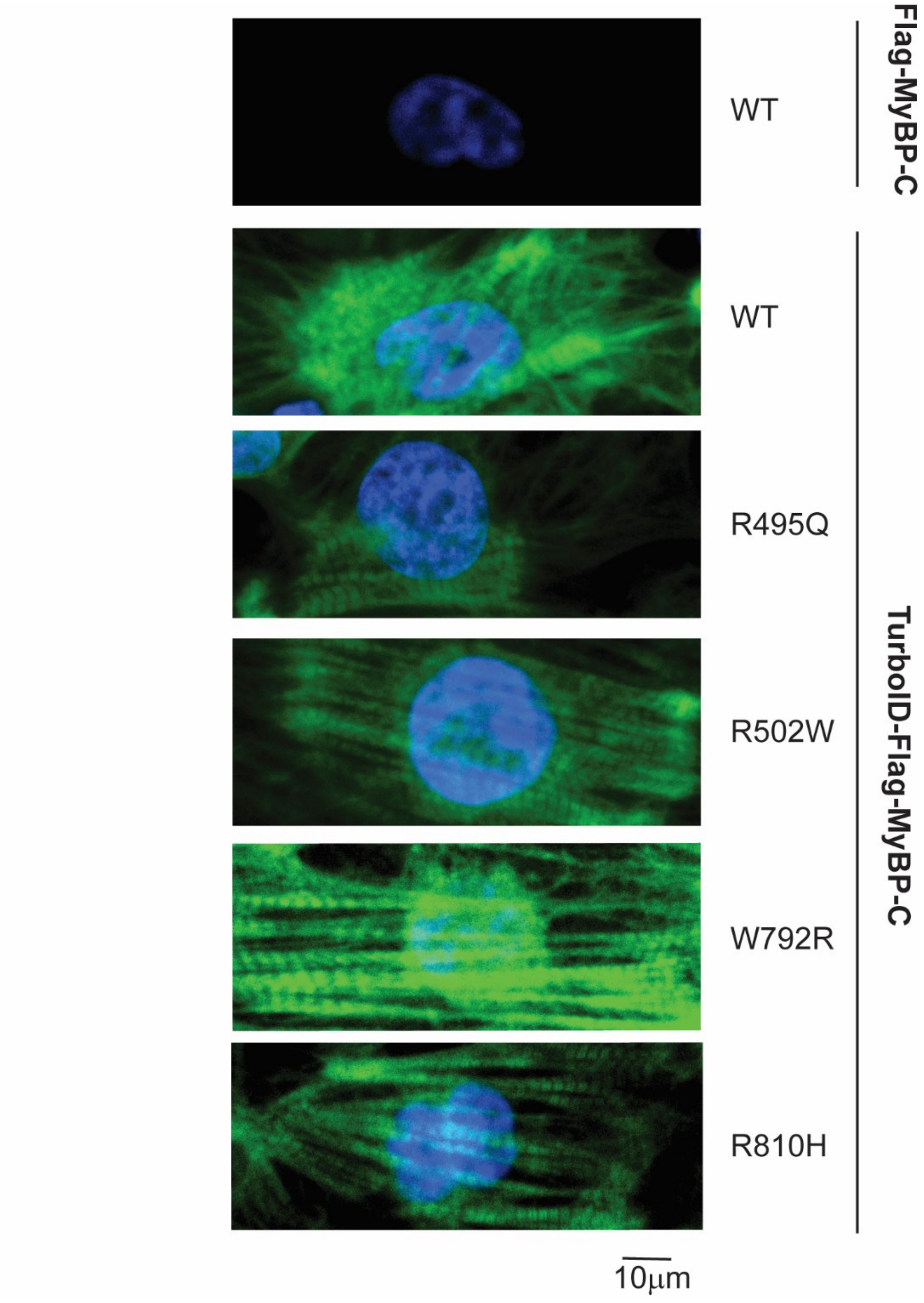


**Supplemental Figure 4: TurboID-flag-MyBP-C protein constructs result in biotinylated proteins within the sarcomere.** The localization of biotinylated proteins after expression of flag-MyBP-C WT (negative control) and TurboID-flag-MyBP-C WT (positive control) as well as Turbo ID-flag MyBP-C P/LP missense variants was visualized by streptavidin-AlexaFlour 488 Immunofluorescence. We observe localization of biotinylated proteins within myofibrils in all TurboID-flag-MyBP-C samples (green). The nucleus is visualized by DAPI staining (blue). A representative image is shown. All samples were tested in biological duplicates with six images collected per biologic replicate.

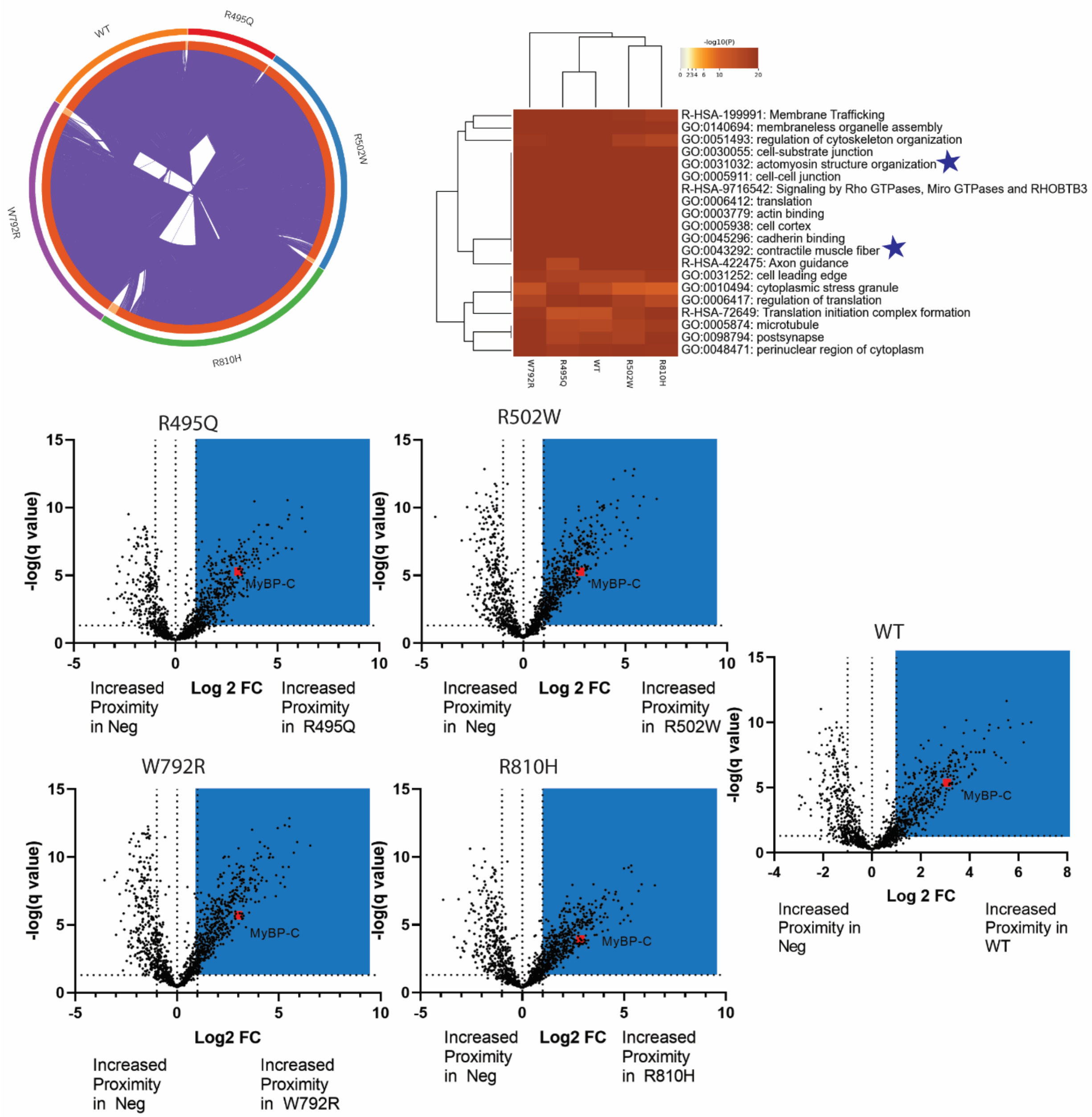


**Supplemental Figure 5: MyBP-C proximity proteins identified by DDA mass spectrometry.** 3,240 MyBP-C proximity proteins (Supplemental Table 9) were identified for WT MyBP-C and/or MyBP-C with P/LP *MYBPC3* missense variants. The circo plot (as described in Supplemental Figure 2), visualizes the significant overlap between the proximity proteins identified across our samples (Top left). A dendrogram (as described in Supplemental Figure 2) demonstrates that gene ontology analysis of the top 500 proximity proteins in all samples identified enrichment in shared pathways across all samples. These pathways include myofibril proteins- actomyosin structure (GO:0031032) and contractile muscle fiber proteins (GO:0043292), denoted by blue stars (top right). MyBP-C proximity proteins can be visualized for each individual variant in the above volcano plots (blue box). Bait protein MyBP-C is labeled with biotin in each of our samples (red square).

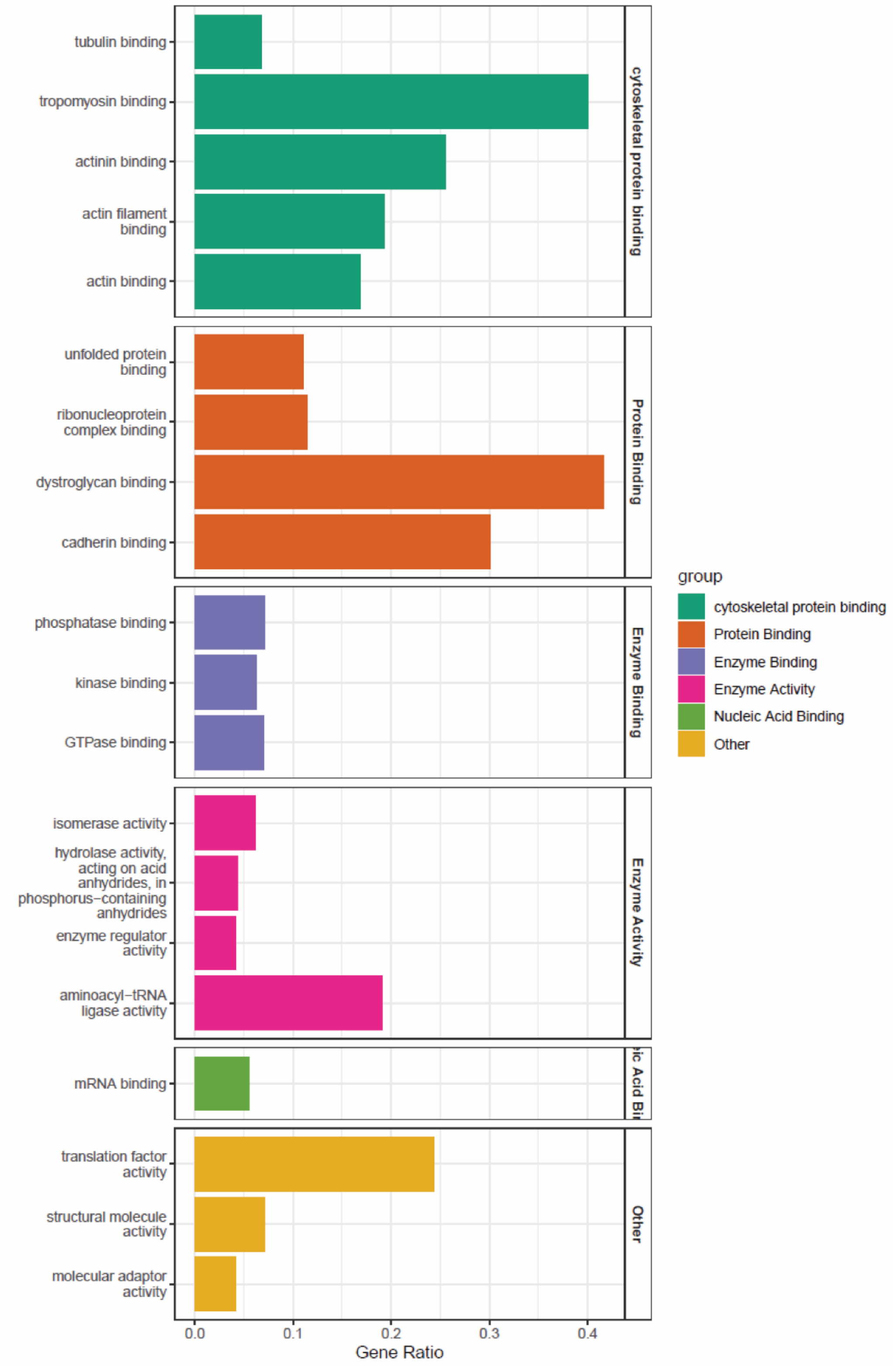


**Supplemental Figure 6: WT MyBP-C Proximity proteins identified by DDA mass spectrometry molecular function gene ontology enrichment analysis.** TOPPGENE was utilized to perform molecular function gene ontology enrichment analysis on the top 500 proximity proteins identified for WT MyBP-C.

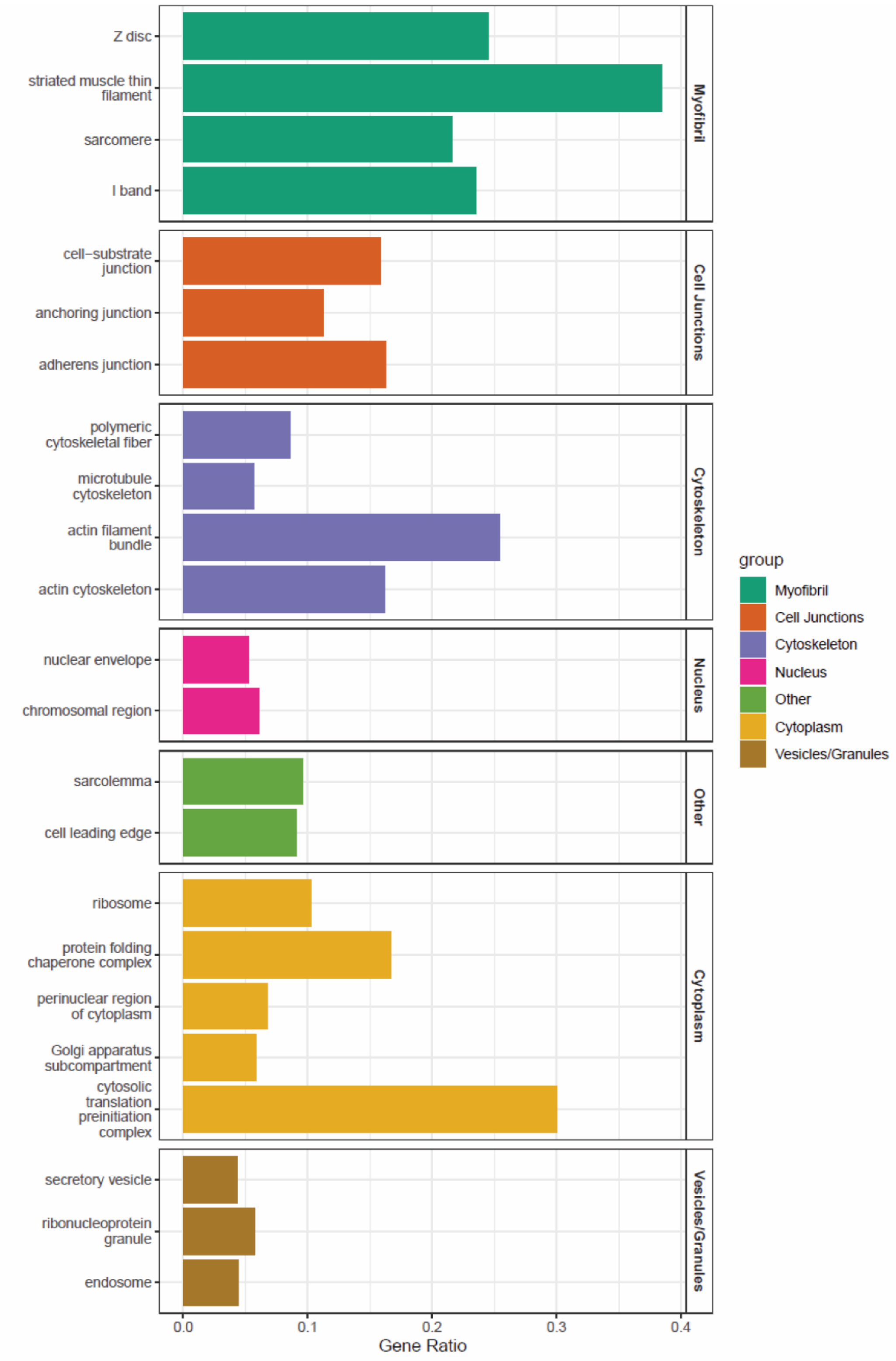


**Supplemental Figure 7: WT MyBP-C Proximity proteins identified by DDA mass spectrometry cellular component gene ontology enrichment analysis.** TOPPGENE was utilized to perform cellular component gene ontology enrichment analysis on the top 500 proximity proteins identified for WT MyBP-C.

**References:**


1. O'Leary TS, Snyder J, Sadayappan S, Day SM, Previs MJ. MYBPC3 truncation mutations enhance actomyosin contractile mechanics in human hypertrophic cardiomyopathy. *J Mol Cell Cardiol*. 2019;127:165-173. doi: 10.1016/j.yjmcc.2018.12.003
2. Glazier AA, Hafeez N, Mellacheruvu D, Basrur V, Nesvizhskii AI, Lee LM, Shao H, Tang V, Yob JM, Gestwicki JE, et al. HSC70 is a chaperone for wild-type and mutant cardiac myosin binding protein C. *JCI Insight*. 2018;3. doi: 10.1172/jci.insight.99319
3. Helms AS, Tang VT, O'Leary TS, Friedline S, Wauchope M, Arora A, Wasserman AH, Smith ED, Lee LM, Wen XW, et al. Effects of MYBPC3 loss-of-function mutations preceding hypertrophic cardiomyopathy. *JCI Insight*. 2020;5. doi: 10.1172/jci.insight.133782
4. Thompson AD, Wagner MJ, Rodriguez J, Malhotra A, Vander Roest S, Lilienthal U, Shao H, Vignesh M, Weber K, Yob JM, et al. An Unbiased Screen Identified the Hsp70-BAG3 Complex as a Regulator of Myosin-Binding Protein C3. *JACC Basic Transl Sci*. 2023;8:1198-1211. doi: 10.1016/j.jacbts.2023.04.009
5. Lian X, Zhang J, Azarin SM, Zhu K, Hazeltine LB, Bao X, Hsiao C, Kamp TJ, Palecek SP. Directed cardiomyocyte differentiation from human pluripotent stem cells by modulating Wnt/beta-catenin signaling under fully defined conditions. *Nat Protoc*. 2013;8:162-175. doi: 10.1038/nprot.2012.150
6. Burridge PW, Holmstrom A, Wu JC. Chemically Defined Culture and Cardiomyocyte Differentiation of Human Pluripotent Stem Cells. *Curr Protoc Hum Genet*. 2015;87:21 23 21-21 23 15. doi: 10.1002/0471142905.hg2103s87
7. Tohyama S, Hattori F, Sano M, Hishiki T, Nagahata Y, Matsuura T, Hashimoto H, Suzuki T, Yamashita H, Satoh Y, et al. Distinct metabolic flow enables large-scale purification of mouse and human pluripotent stem cell-derived cardiomyocytes. *Cell Stem Cell*. 2013;12:127-137. doi: 10.1016/j.stem.2012.09.013
8. Helms AS, Thompson AD, Glazier AA, Hafeez N, Kabani S, Rodriguez J, Yob JM, Woolcock H, Mazzarotto F, Lakdawala NK, et al. Spatial and Functional Distribution of MYBPC3 Pathogenic Variants and Clinical Outcomes in Patients With Hypertrophic Cardiomyopathy. *Circ Genom Precis Med*. 2020;13:396-405. doi: 10.1161/CIRCGEN.120.002929
9. Zhou Y, Zhou B, Pache L, Chang M, Khodabakhshi AH, Tanaseichuk O, Benner C, Chanda SK. Metascape provides a biologist-oriented resource for the analysis of systems-level datasets. *Nat Commun*. 2019;10:1523. doi: 10.1038/s41467-019-09234-6
10. Chen J, Bardes EE, Aronow BJ, Jegga AG. ToppGene Suite for gene list enrichment analysis and candidate gene prioritization. *Nucleic Acids Res*. 2009;37:W305-311. doi: 10.1093/nar/gkp427
11. Helms AS, Davis FM, Coleman D, Bartolone SN, Glazier AA, Pagani F, Yob JM, Sadayappan S, Pedersen E, Lyons R, et al. Sarcomere mutation-specific expression patterns in human hypertrophic cardiomyopathy. *Circ Cardiovasc Genet*. 2014;7:434-443. doi: 10.1161/CIRCGENETICS.113.000448
